# Quantitative comparison of fluorescent reporters by FCS excitation scan

**DOI:** 10.64898/2026.04.04.716477

**Authors:** Falk Schneider, Le A. Trinh, Scott E. Fraser

**Affiliations:** Translational Imaging Center, Michelson Center for Convergent Bioscience, University of Southern California, Los Angeles, CA, 90089, USA; Molecular and Computational Biology, University of Southern California, Los Angeles, CA, 90089, USA; Directorate of Biomedical Sciences, Warwick Medical School, University of Warwick, Coventry CV4 8UW, UK; Biohub, 1180 Main, Redwood City, CA 94063, USA

## Abstract

Fluorescent reporters such as fluorescent proteins or chemigenetic indicators are indispensable tools for studying biological processes using light microscopy. Choosing an appropriate fluorescent tag is a crucial step in experimental design not only for imaging but also for quantitative measurements such as fluorescence fluctuation spectroscopy. Two key parameters should be considered: Fluorescent brightness and photo-bleaching. Change to fluorescence intensity due to photobleaching is relatively easy to assess in different biological environments, while brightness is more elusive. Here, we develop and employ a fluorescence correlation spectroscopy (FCS) based excitation scan assay that determines fluorescent protein performance and validate it in tissue culture and zebrafish embryos. We employ our FCS pipeline to compare a set of 10 established fluorescent proteins as well as HALO and SNAP tags for both cellular imaging and measurements of diffusion dynamics with FCS. We show that mNeonGreen outperforms mEGFP in tissue culture and zebrafish embryos. We also compare StayGold variants against other green fluorescent proteins and chemigenetic reporters in tissue culture. Overall, we present a broadly applicable approach for determining fluorescent reporter brightness in the living system of interest.

## Introduction

Fluorescent tags are indispensable tools in fluorescence microscopy, enabling the visualisation of specific molecular mechanisms in living systems. Among these tags, the introduction of fluorescent proteins (FPs) has basically revolutionised cellular imaging since the discovery, cloning, and recombinant expression of the Green Fluorescent Protein (GFP) from *Aequorea victoria*^1,2^. Over the decades, discovery and extensive engineering of FPs and biosensors has enhanced their photostability, brightness, excitation/emission wavelengths, and fluorescence lifetimes, broadening their applications in the life sciences^3,4^. Particularly exciting has been the introduction of the photo-stable StayGold protein^5^ and its monomeric variants^6–8^.

Alongside FPs, chemigenetic labelling tools have emerged as versatile alternatives. Initial systems such as FLAsH/ReAsH peptides bind non-covalently to biarsenical fluorophores^9^. Enzymatic systems such as HALO-^10^, SNAP-^11^, or CLIP-tags^12^ that use self-labelling to covalently attach organic fluorophores, offering precise control over fluorescence properties through the choice of dye. Chemigenetic tools are particularly suited to advanced microscopy techniques, such as single-molecule localisation microscopy (SMLM) and stimulated emission depletion (STED) microscopy, which demand specific photophysical properties from their imaging probes^13,14^.

While the number of available fluorescent markers is steadily increasing, there are few objective methods to quantitatively compare them. Thus, selecting the optimal tag for a given application remains challenging and can make or break an experiment. Key parameters include brightness, i.e., number of photons per fluorophore per time, photostability, fluorescence lifetime, quantum yield, extinction coefficient, and oligomeric state^15^. Such properties must be rigorously characterised in the context of their biological environment, as factors such as pH, salinity, and cellular milieu can significantly affect fluorescence behaviour^16–18^. It should be noted that rarely a single fluorescent reporter fits every experimental need

The brightness of a fluorophore essentially determines how many photons can be recorded and therefore critically contributes to the signal-to-noise ratio (SNR) in an imaging experiment. Traditionally, brightness is estimated using *in vitro* assays and recombinant proteins, with extinction coefficients and quantum yields determined spectrophotometrically^19,20^. While these measurements provide valuable benchmarks, they often fail to capture the complexities of cellular environments. Assays based on polycistronic expression systems elegantly address this by co-expressing two FPs to match expression levels, enabling relative brightness comparisons^21^. However, these methods rely on relative intensity measurements that can be difficult to interpret given varying quantum efficiencies of detectors at different wavelength.

Fluorescence correlation spectroscopy (FCS) circumvents these limitations by directly measuring molecular brightness without relying on an internal standard^22^. FCS quantifies diffusion dynamics, concentration, and molecular brightness of fluorescently labelled molecules by analysing fluorescence fluctuations recorded at a single spot on a confocal microscope^23–25^. The brightness metric is particularly useful for inferring oligomeric states^26^ and utilised in other fluorescence fluctuation spectroscopy techniques such as Number&Brightness (N&B) analysis^27,28^. In FCS, the SNR depends on the molecular brightness, diffusion coefficient, and the acquisition time^24,29^. While diffusion dynamics are intrinsic to the system under study and acquisition times are often limited by biological constraints, brightness is the key parameter that can be changed by the choice of fluorophore.

In this study, we present a method to quantitatively determine and compare the brightness of fluorescent probes in different biological environments. We employ an FCS excitation scan measuring brightness and diffusion dynamics against varying laser power. We demonstrate the approach in two distinct systems: tissue culture using human embryonic kidney (HEK) 293T cells and *in vivo* imaging of living zebrafish embryos in hindbrain tissue. By correlating brightness with other photophysical properties across varying excitation powers, this method provides a robust framework for evaluating fluorescent probe performance under biologically relevant conditions.

The principle of acquiring FCS data at different excitation powers is illustrated schematically in **Figure 1a**. Intensity fluctuations from fluorophores diffusing through the confocal observation volume are recorded and used to calculate the FCS autocorrelation curves. From fitting these, diffusion dynamics and brightness can be extracted. Approaches like this have been leveraged previously to demonstrate fluorescence reporter performance *in vitro* or in cell culture^24,30,31^ but have not been applied *in vivo*. For the same probe, higher incident laser power produces higher fluorescence signals, improving the SNR of the FCS acquisition. While in focus, the fluorophore emits photons at a rate that depends on its brightness, measured as counts per molecule (cpm), and the incident laser power. The cpm are derived from the fitted amplitude of the FCS curves and the first moment of the fluorescence intensity^24,26^.

**Figure 1:**
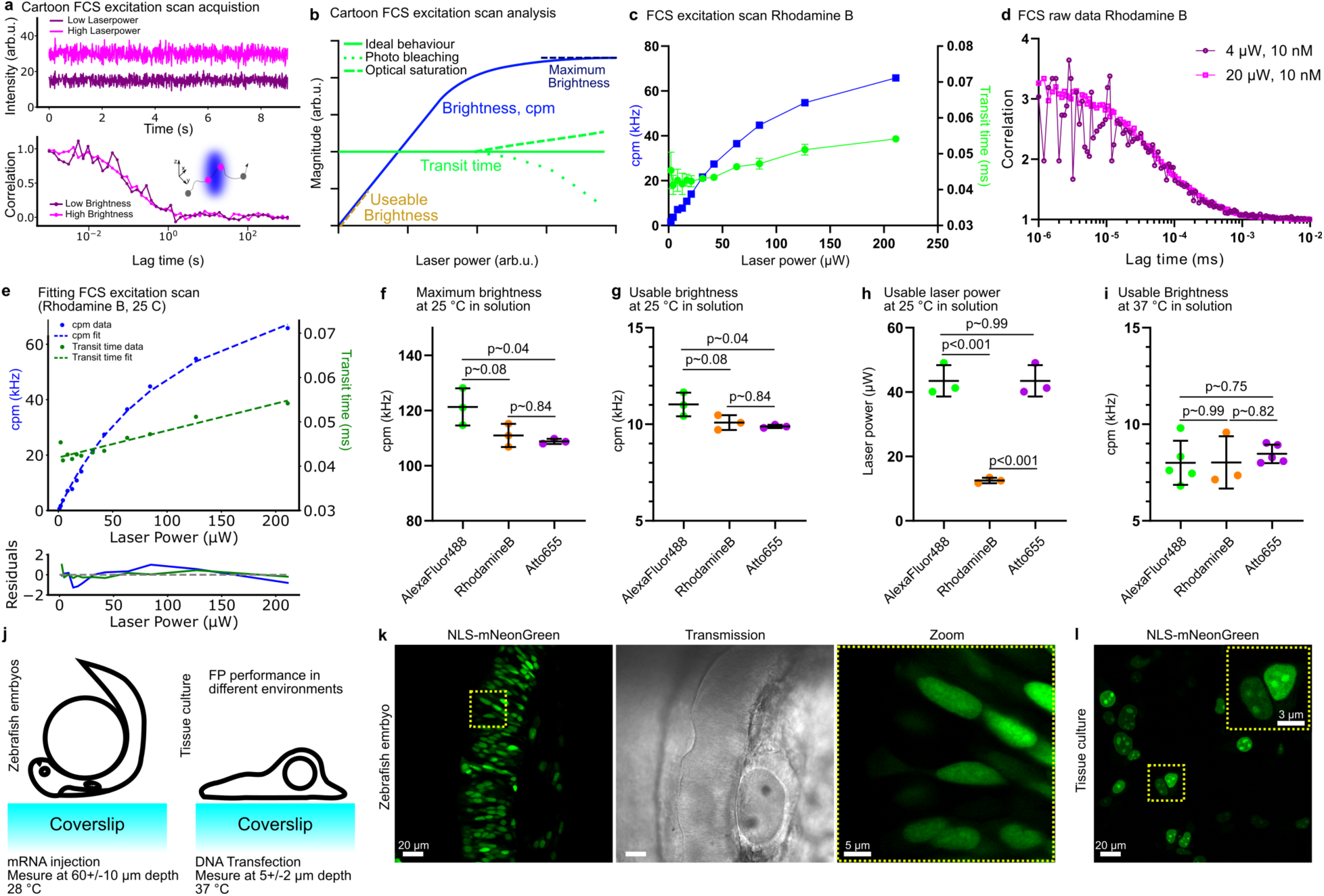
FCS excitation scan to measure and compare fluorescent probe brightness. **a** Cartoon of FCS excitation scan acquisition at two different laser powers. Top panel change in intensity, bottom panel change to noise in the FCS curve due to change in brightness caused by different excitation laser powers (lower laser power results in lower brightness). **b** Cartoon of the FCS excitation scan analysis. Transit time and brightness are measured against laser power. Ideally transit time does not chance with laser power, photo-bleaching or optical saturation cause changes. Brightness saturates at high laser power. FCS measurements should be performed in the linear regime of the curve (compare **Supplementary Figure S1**). **c** FCS excitation scan of Rhodamine B in water at room temperature. **d** FCS curves of Rhodamine B in water at room temperature at different excitation powers. **e** Fitting of the FCS excitation scan data for Rhodamine B to extract maximum brightness, usable brightness, and usable laser power. **f-h** FCS excitation scan results for AlexaFluor 488, Rhodamine B and Atto655 in solution at room temperature (25 °C). Maximum brightness (**f**), usable brightness (**g**), and usable laser power (**h**). **i** Usable laser power of AlexaFluor 488, Rhodamine B, and Atto655 at 37° C. All adjusted p-values were obtained from one-way ANOVA with Tukey’s correction for multiple comparisons. **j** Cartoon of experimental setup for FCS excitation scans in zebrafish (left) and HEK293T cells (right). **k** Confocal image of zebrafish hindbrain, transmission image and zoom in. Nuclei labelled with NLS-mNeonGreen. l Confocal image of HEK293T cells transfected with NLS-mNeonGreen, inlet shows zoom in.

In an FCS excitation scan (illustrated in **Figure 1a,b** and **Supplementary Figure S1**), FCS fitting results are plotted at varying laser powers. Under ideal conditions, the molecular transit time (the average time molecules take to traverse the observation volume) is independent of laser power. However, photophysical phenomena such as photobleaching and optical saturation can cause deviations from this behaviour (**Figure 1b** and **Supplementary Figure S1c,d**)^32,33^. For instance, photobleaching causes the fluorophore to stop emitting photons before leaving the observation volume, which decreases the effective transit time. The brightness of a fluorophore ideally increases linearly with laser power which holds at low excitation powers. However, in practice it follows a saturation curve as the excitation approaches the saturation regime where emission rate is limited by cycling through ground and excited state. To ensure quantitative FCS measurements, it is important to operate in the linear regime of fluorescence excitation, where brightness scales linearly with laser power^32,34–36^. The brightness determined in this regime provides a robust, quantifiable measure of fluorescent probe performance for FCS and imaging applications. We refer to this as ‘useable’ brightness in contrast to ‘maximum’ brightness obtained under saturating conditions (schematically shown in **Figure 1b**). Here, we employ the FCS excitation scan to determine the brightness of 10 commonly used FPs in tissue culture and in vivo and identify mNeonGreen as the best performing reporter in both systems. Furthermore, we test chemigenetic reporters as well as the recently developed monomeric StayGold versions in tissue culture.

## Results

First, we demonstrate the versatility and applicability of the FCS excitation scan approach by comparing three fluorescent dyes: Alexa Fluor 488 (AF488), Rhodamine B (RhB), and Atto655 in aqueous solution. These widely used, bright, and well-characterised probes provide demonstration for the brightness values expected at the upper end of the performance spectrum. **Figure 1c** shows an FCS excitation scan for RhB in water at room temperature. The increase in cpm with laser power saturates. The transit time is initially constant, then increases revealing optical saturation at higher laser powers (compare schematic in **Figure 1b**). Transit time values at low laser power (∼ 1 μW, cpm < 1 kHz) show large standard deviation. The influence of the incident laser power (and resulting brightness) is illustrated in **Figure 1d** with higher SNR (less spread) of the correlation curve for higher power, higher brightness.

Useable brightness and the saturation curve are molecular properties. To compare different fluorophores, we propose to fit the FCS excitation scan data for cpm and transit time (example for RhB in **Figure 1e**). The cpm curve is best fit by a saturation model and the transit times by a linear model. The resulting fitting parameters deliver the maximum brightness, brightness at saturation, and the usable brightness defined as 10% of the inflection point of the saturation curve (illustration in **Figure 1b**). In the majority of cases, the saturation drives the value for the usable brightness. Yet, in some cases photo-bleaching occurs before the 10% inflection in brightness point. In these cases, we use the point where the transit time dropped by less than 20%. Note that the linear model for the transit time and the saturation model for the brightness are simplifications. The exact photophysical processes and dependences can be rather complex. However, our simple empirical approach describes the data well, provides a means for straightforward analyses, and allows comparison of data from multiple repetitions, fluorophores, and conditions. The fitting results from the FCS excitation scans on AF488, RhB, and Atto655 are summarised in **Figure 1f-I** (**Supplementary Figure S2** shows examples of FCS curves and fits for all three dyes at different excitation powers). The maximum brightness (**Figure 1f**) scales with the useable brightness (**Figure 1g**). In both cases AF488 shows the highest value (usable brightness: 11.0+/-0.6 kHz) and RhB and Atto655 comparable values (10.1+/-0.4 kHz and 9.9+/-0.1 kHz, respectively). Interestingly, the laser power required to reach the useable brightness is less than half for RhB (12.5+/-0.8 μW) compared to the other dyes (43.5+/-4.9 μW for AF488 and 43.5+/-4.9 μW for Atto655). These features are important since in a biological experiment the sample will be exposed to a much lower light dose to reach the same SNR. Furthermore, most quantitative imaging approaches rely on linear increase of fluorescence with number of fluorophores which is not given for measurements performed at higher excitation powers. The FCS excitation scan and the presented fitting strategy thus allow comparing different fluorophores for their brightness and useable laser power. This allows comparison at different conditions, for example, at different temperatures where it is already known that different dyes can have different photon outputs^37^. Correspondingly, we find a decrease in useable brightness for all dyes (**Figure 1i**; examples of excitation scan fits are shown in **Supplementary Figure S3**) rendering the brightness of the three dyes at 37 °C comparable to each other. These results highlight the importance of testing and validating fluorescent reporters under different experimental conditions.

Next, we used this pipeline to compare the performance of different fluorescent proteins for FCS in cell culture and zebrafish embryos with the main differences between the two system being imaging depth and temperature: about 60 μm at 28 °C for the zebrafish embryos and 5 μm at 37 °C for the HEK cells (**Figure 1j**). We employed a dual-purpose expression construct supporting *in vitro* transcription to produce mRNA for zebrafish embryo injection (SP6 promotor) and transfection into mammalian cells (CMV promotor, see Materials and Methods). **Figure 1k,j** shows expression of nuclear localisation signal (NLS)-tagged mNeonGreen in the hindbrain of ∼20 hours post-fertilization (hpf) zebrafish embryos and in HEK293T cells. The NLS-tag was chosen to demonstrate feasibility of measurements in a specific cellular compartment, i.e. in the nucleus within complex samples.

We chose a set of 10 monomeric FPs across the visible spectrum based on their reported brightness: mCerulean3, mTurquoise2, mNeonGreen, mEGFP, mKOkappa, stagRFP, mSCarlet, mCherry, emiRFP670, and miRFP670nano3. All of the FPs were fused to an N-terminal NLS and expressed in both systems (**Figure 2a,b**). FCS excitation scans were performed across multiple nuclei and embryos with experiments independently repeated (see Materials and Methods; see **Supplementary Figure S4,S5** for representative FCS data). Despite potential differences between imaging depth or exact mounting orientation, experiments were reproducible in independent biological repeats (examples for mNeonGreen and mEGFP in zebrafish embryos shown in **Figure 2c,d**; HEK cells: **Figure 2e,f**). From fitting the FCS excitation scan data, we extracted usable brightness and usable laser power for the 10 FPs in cell culture and live embryo model systems (**Figure 2g-h**; representative excitation scan fits in **Supplementary Figure S6**) across the blue, green, orange and red wavelength ranges.

**Figure 2:**
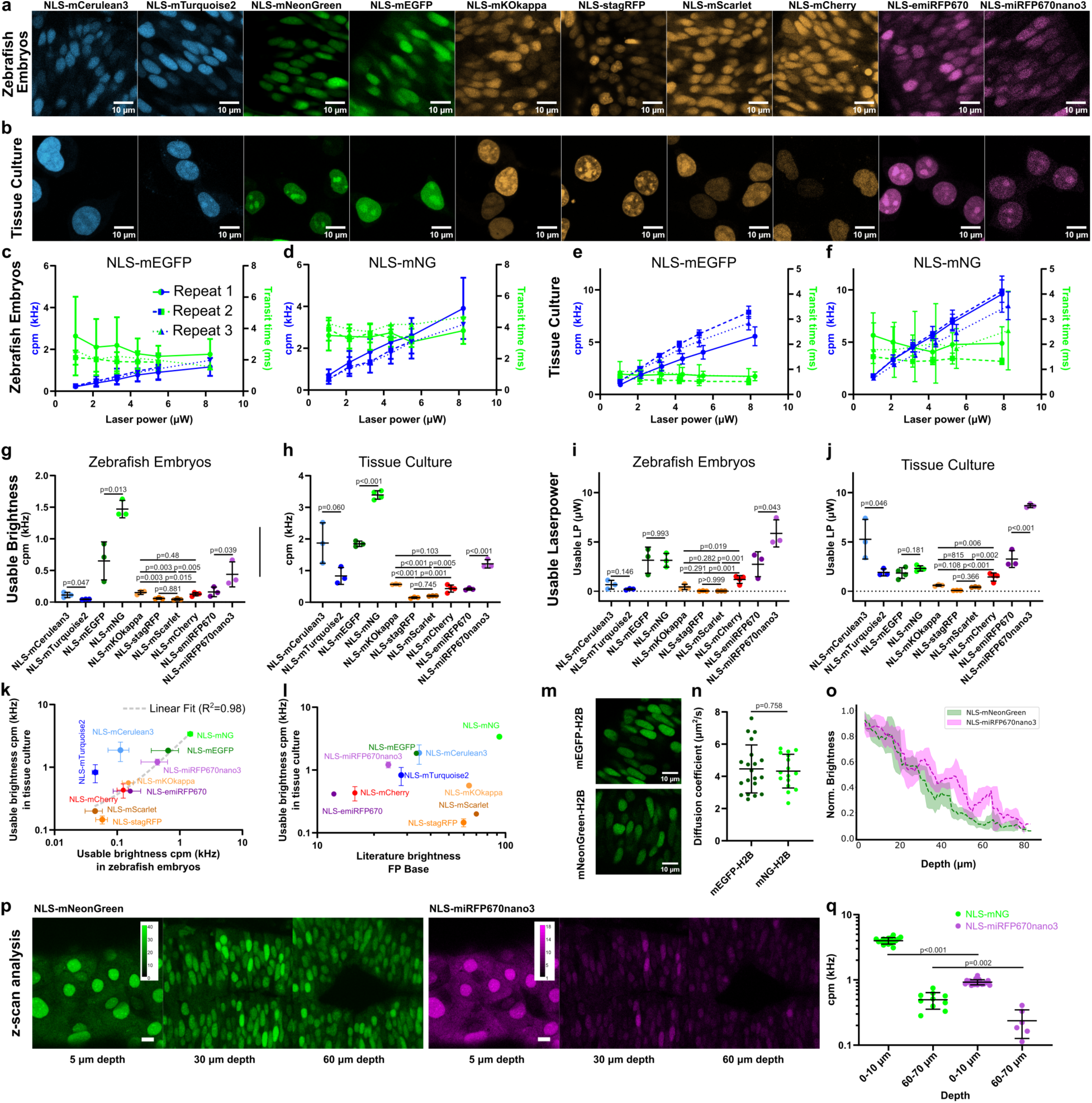
Brightness and performance of 10 fluorescent proteins in zebrafish embryos and tissue culture. **a,b** confocal images of cells in the zebrafish hindbrain (∼50 um depth, **a**) and HEK293T cells (**b**) expressing ten nuclear localized fluorescent proteins. **c,d** FCS excitation scan of NLS-mEGFP and NLS-mNG expressed in cells in zebrafish hindbrain at 28° C. Repeats are three independent experiments. For every independent repeat 3-5 FCS curves at every laser power have been acquired in at least 3 nuclei for at least 6 embryos. **e,f** FCS excitation scan of NLS-mEGFP and NLS-mNG in HEK293T cells at 37° C. For every independent repeat at 3-5 FCS curves at every laser power have been acquired in at least 10 nuclei. **g,h** Usable brightness obtained from the FCS excitation scans in zebrafish embryo hindbrain and tissue culture. **i,j** Usable laser power obtained from the FCS excitation scans in zebrafish embryo hindbrain and tissue culture. Both usable brightness and laser power are outputs from fitting the FCS excitation scans. Every dot refers to one repeat (independent experiment). Lines are average values and bars standard deviation. Adjusted p-values were obtained from student’s t-test for blue, green and red FPs. Adjusted p-Values for orange FPs were obtained from one-way ANOVA with Tukey’s correction for multiple comparisons. For statistical comparison of FPs across different excitation wavelength see **Supplementary Tables T1-T4**. **k** Scatter plot between usable brightness in zebrafish embryo hindbrain and HEK293T cell culture. A linear fit was applied to the data (excluding the blue excitable proteins NLS-mTurquoise2 and NLS-mCerulean3). **l** Scatter plot between brightness data from the literature (obtained from FP base) and the usable brightness from the FCS excitation scans in HEK293T cells. **m** Confocal images of nuclei expressing mEGFP-H2B and mNeonGreen-H2B in the zebrafish hindbrain. **n** Diffusion coefficients calculated from FCS measurements on mEGFP-H2B and mNeonGreen-H2B in zebrafish hindbrains. Every dot represents one nucleus and averaging over 3-5 FCS curves. Data were acquired on 6 individual zebrafish embryos. Horizontal line represents mean, bars are standard deviation. **o** Normalized brightness of NLS-mNeonGreen and NLS-miRFP670nano3 as a function of imaging depth into the zebrafish hindbrain. Line represents average and shade standard deviation calculated from moving average. **p** Confocal images of NLS-mNeonGreen and NLS-miRFP670nano3 labelled nuclei at different depth in the zebrafish hindbrain. **q** Brightness (cpm) of NLS-mNeonGreen and NLS-mirFP670nano3 in nuclei of zebrafish embryos at 0-10 and 60-70 um depth. Every dot represents one nuclei averaging over 5 FCS curves. Error bars are standard deviation. Adjusted p-value from pairwise comparison at different depth were obtained using student’s t-test.

In the green emission range, mNeonGreen displays noticeably the highest usable brightness among the compared proteins with 1.5+/-0.1 kHz in the embryos and 3.4+/-0.1 kHz in the HEK cells being 2.3 times (zebrafish embryos) and 1.8 times (HEK cells) brighter than mEGFP. Intriguingly, the usable laser powers for mNeonGreen and mEGFP are similar, indicating that the same amount of incident light leads to higher SNR for mNeonGreen. In the blue wavelength range, mCerulean3 outperforms mTourquise2 but with higher laser power required. Furthermore, the blue FPs show very poor excitation and brightness in the zebrafish embryo (at depth). In the orange emission regime, mKOkappa and mCherry are good choices, with mKOkappa having slightly higher brightness (1.3 times brighter as compared to mCherry in both model systems). Finally, miRFP670nano3 showed the highest brightness in the red regime. The usable brightness was higher compared to, for example, mCherry or mScarlet, however, to reach usable brightness the largest light doses in our comparison had to be used. Yet, this is still less than 10 μW at the sample which is commonly used for FCS studies on dyes in solution or on membranes^13^.

The usable brightness data in zebrafish embryos and tissue culture model show a good correlation excluding mCerulean3 and mTurquoise2 (**Figure 2k**, R^2^=0.98). These blue-excitable FPs appear dimmer in the embryos as expected due to the higher absorbance and scattering of blue shifted light^38^. The correlation for the other 8 FPs in both models indicates that expression and FCS performance in HEK cells can be a good predictor for performance in zebrafish embryos and potentially other model systems. When compared to literature values from FPbase.org^39^, the usable brightness data in the HEK cells shows only a partial correlation (**Figure 2l**). While the brightness from FCS (cpm) and the spectrophotometrically determined brightness in the literature (FPbase.org) refer not to the exact same property, they can be used as a starting point to choose probes. Clear outliers in this scatter plot are stagRFP and mScarlet. These FPs show pronounced photo-bleaching and a large contribution of dark state kinetics to their raw autocorrelation data (**Supplementary Figure S7**), providing an explanation for this divergence. These deviations represent a distinguishing feature of these two candidates from the other FPs contributing to the lower brightness as expected from literature (**Figure 2l**).

To demonstrate the importance and impact of choosing the right fluorescent protein for FCS experiments, we chose to measure the diffusion coefficient of histone H2B N-terminally tagged with mEGFP and mNeonGreen in nuclei of the zebrafish hindbrain (**Figure 2m**). While the average diffusion coefficient is almost identical as driven by the diffusion dynamics of H2B (D(mEGFP-H2B)=4.4+/-1.5 μm^2^/s and D(mNG-H2B)=4.3+/-1.0 μm^2^/s), the spread, standard deviation, of the data is larger for the mEGFP tagged version implying that less measurements are required to gain the same level of confidence with the mNeonGreen than mEGFP (mNeonGreen higher brightness is maintained in the H2B fusion constructs, **Supplementary Figure S8**).

To measure diffusion dynamics at depth, we compared the performance of the brightest fluorescent protein tested thus far, mNeonGreen, to a less bright but red-shifted FP with the expectation that red-shifted light is less scattered and also less absorbed by the tissue and hence, red-shifted FPs may outperform brighter FPs for FCS experiments. We performed FCS measurements and confocal imaging at different depth and assessed diffusion dynamics and brightness (**Figure 2o,p**). The normalised brightness drops with depth noticeably slower for the red-shifted miRFP670nano3 as compared to mNeonGreen. Nevertheless, the FCS data quality is determined by the absolute value of the brightness, the number of photons per molecule reaching the detector matters. While we see a stronger drop in brightness for mNeonGreen, the absolute value at 60-70 μm depth of the cpm is still higher as compared to miRFP670nano3 (0.49+/-0.23 versus 0.23+/- 0.11 kHz respectively, **Figure 2q**). This implies that it is advantageous to use the brightest available probe when imaging deeper, even if there is more scattering of the emitted photons.

The measurements at different depth clearly illustrate the importance of high initial brightness. Chemigenetic labels employ organic fluorophores and are considered to be brighter and more photostable as compared to fluorescent proteins. We applied FCS excitation scan to validate this assumption in the tissue culture model. As we found good correlation between the green, orange and red fluorescent protein brightness in the two model systems (**Figure 2k**), measurements in HEK cells can serve as a proxy for performance in the zebrafish. The most commonly used chemigenetic labelling systems are HALO and SNAP. We chose two popular, commercially available, green and red fluorescent ligands, SNAP-Cell 505 Star, Oregon Green (HALO ligand), Janelia Fluor 646 (HALO ligand), and SNAP-Cell 647SiR, due to their widespread use and commercial availability. First, we compared usable brightness and laser power at 37 °C for the free ligands in aqueous solution (**Figure 3a,b; Supplementary Figure S9**). Oregon Green shows poor performance in FCS being about ten times dimmer as compared to other dyes due to quickly reaching optical saturation. The usable brightness for SNAP-Cell 505-Star (11.4+/-1.9 kHz), Janelia Fluor 646 (8.6+/-1.0 kHz), and SNAP-Cell 647-SiR (7.6+/-0.9 kHz) are on the same order as the organic fluorophores at 37 °C shown in **Figure 1i**.

**Figure 3:**
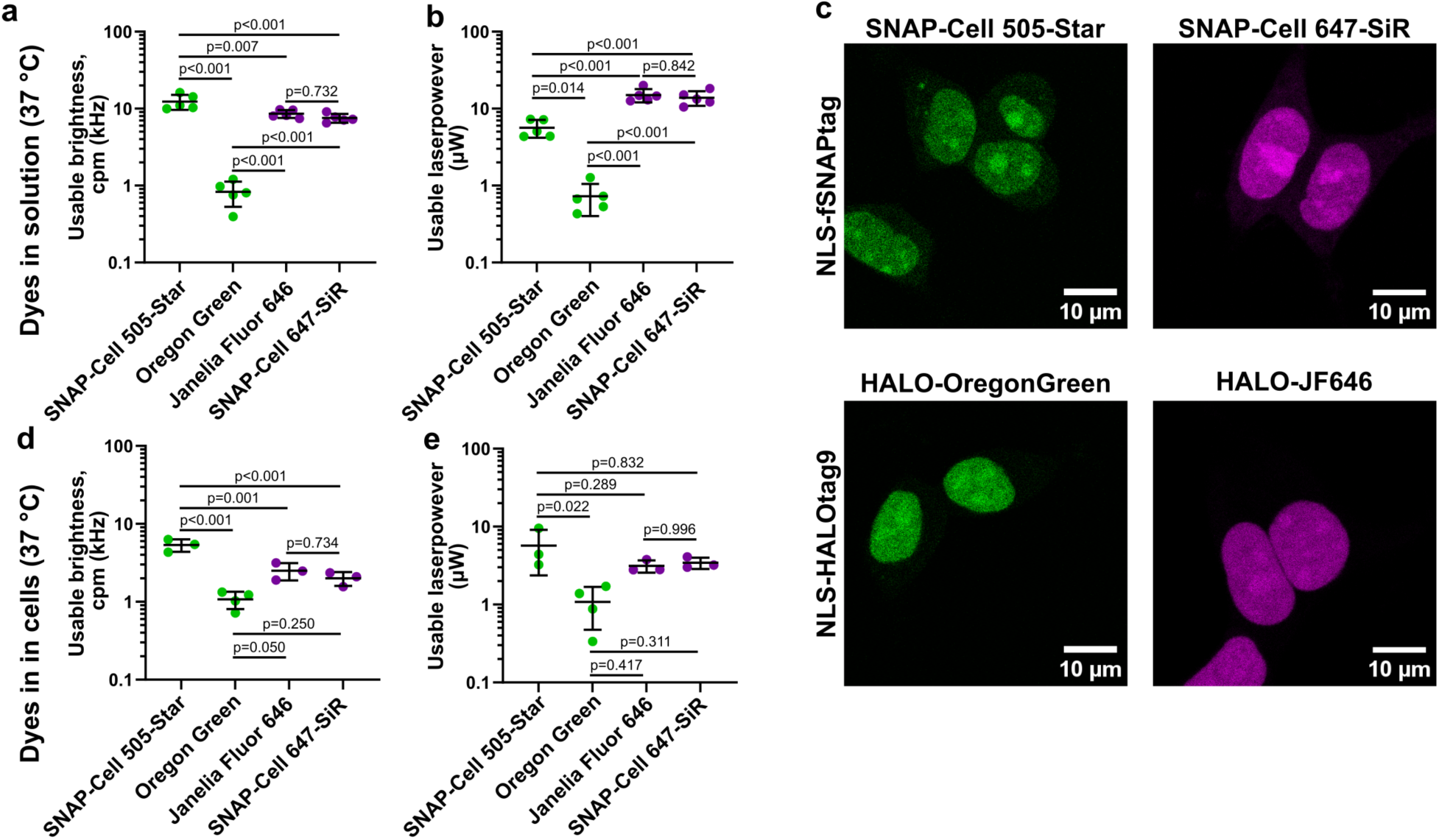
Brightness and performance of the chemigenetic labelling systems HALOtag9 and fSNAPtag with common ligands. **a** Usable brightness and **b** usable laser power from FCS excitation scans of SNAP-Cell 505-Star, Oregon Green HALO ligand, Janelia Fluor 646 HALO ligand, and SNAP-Cell 647-SiR in solution at 37° C. **c** Confocal images of HEK293T cells expressing HALOtag9 or fSNAPtag and labelled with the indicated ligand. **d** Usable brightness and **e** usable laser power from FCS excitation scans of NLS-HALOtag9 and NLS-fSNAPtag expressed in HEK293T cells and labelled with SNAP-Cell 505-Star, Oregon Green HALO ligand, Janelia Fluor 646 HALO ligand, and SNAP-Cell 647-SiR at 37° C. Every dot is representative of one experiment (average). Horizontal bars are mean values and error bars are standard deviations. Adjusted p-values were obtained from one-way ANOVA with Tukey’s correction for multiple comparisons.

To assess brightness and FCS performance of the dyes in a more physiological environment, we used fast SNAPtag (fSNAP)^40^ and HALOtag9^41^. Similarly to the FPs, they were expressed in HEK cells fused to an NLS-tag (**Figure 3c**). The usable brightness and usable laser power dropped slightly for all ligands, except for Oregon Green (**Figure 3d,e**). The useable brightness of NLS-HALOtag9-Janelia Fluor 646 (2.5+/-0.6 kHz) and NLS-fSNAPtag-SNAP-Cell 647-SiR (2.0+/-0.4 kHz) are higher than the usable brightness of miRFP670nano3 (1.2+/-0.1, **Figure 2h**) and require substantially less excitation power (3.1+/-0.6 μW and 3.4+/-0.6 μW, respectively, compared to 8.7+/-0.2 μW for miRFP670nano3). NLS-fSNAPtag-SNAP-Cell-505-Star shows a brightness of 5.4+/-0.9 kHz and is thus brighter than NLS-mNeonGreen (3.4+/-0.1 kHz) in the HEK cells. These measurements confirm the overall higher useable brightness for some of the organic dyes. However, we were not able to wash all free dye out and had to fit the FCS curves with a two-component model potentially complicating biological analysis (**Supplementary Figure S10**). The presence of free dye in the cells detected in the FCS experiments may cause complications in interpreting the data.

Finally, we applied FCS excitation scan to more recently developed green fluorescent proteins based on the dimeric StayGold^5^. Specifically, we tested four different variants and follow the naming convention from FPbase.org: mSG, mSG2^6^, mBaoJin^8^ and SG-E138D^7^. All FPs were expressed as previously with an N-terminal NLS-tag in HEK cells (**Figure 4a**). We did not observe any obvious differences in levels of expression. Maximum brightness of all FPs is above 90 kHz (**Figure 4b**) and thus on the order of the organic dyes in **Figure 1f**. Usable brightness values (**Figure 4c**) in the FCS excitation assays for mSG and mSG2 surpass the values for the organic dyes in **Figure 1f,g** and **Figure 3d** with values of 14.9+/-2.0 kHz and 14.6+/-0.9 kHz respectively (see representative FCS data and excitation scans in **Supplementary Figure S11, S12**). The other two StayGold variants are about 0.6 times dimmer and show similar values for useable brightness as compared to the dyes in **Figure 1g**. Noticeably, the useable laser powers are all well below 10 μW and thus only a quarter of the laser power required for AF488 to reach the same useable brightness. Finally, a distinguishing feature of the StayGold variants is the photo-stability as compared to other fluorescent proteins. Comparison with NLS-mNeonGreen in the FCS excitation scan assay reveals clear decrease in transit time of NLS-mNeonGreen but not the StayGold variants over the course of investigated laser powers (**Figure 4e**) highlighting the advantage in long-term imaging for the StayGold variants.

**Figure 4:**
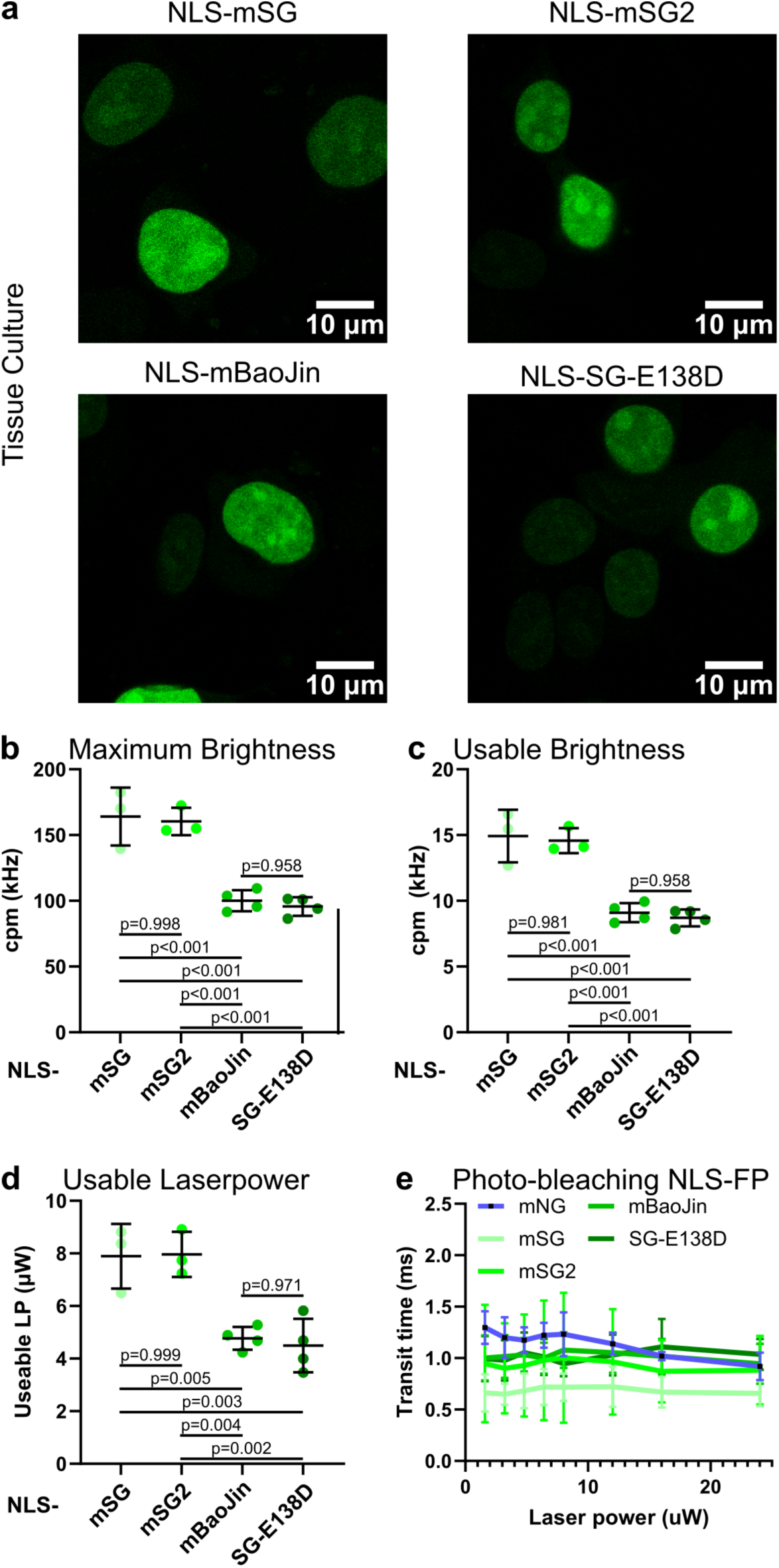
Performance and brightness of monomeric StayGold variants in FCS excitation scan. **a** Confocal images of HEK293T cells expressing NLS-mSG, NLSmSG2, NLS-mBaojin, and NLS-SG-E138D. FCS excitation scans were performed at 37° C. Results are shown as Maximum brightness (**b**), usable Brightness (**c**) and usable laser power (**d**). Every dot represents the result from fitting one independent repetition of an FCS excitation scan. Horizontal line represents average and bars are standard deviation. Adjusted p-values were obtained from one-way ANOVA with Tukey’s correction for multiple comparisons. **e** Dependence of the transit time of the monomeric StayGold variants (green shades) and mNeonGreen (mNG, blue) on incident excitation laser power. Values are mean of 10 nuclei per laser power (3-5 FCS curves per nuclei per laser power). Errors are standard deviations.

## Discussion

Choosing the right fluorophore for an experiment is an important may determine its outcome. This study highlights the importance of brightness, i.e., useable photon flux as selection criterion. We present our FCS-excitation scan pipeline to compare brightness of fluorescent probes for their use for live imaging *in vivo*. This method has been used to compare various FPs in two biological systems, the zebrafish embryo and HEK293T cells, and to compare organic fluorophores to FPs. In addition to brightness, we consider the useable laser power as an important parameter because it determines the amount of light a biological sample has to be exposed to in order to obtain the useable brightness. We find that the high brightness of mNeonGreen and the recently developed StayGold variants, specifically, mSG2, represent distinguishing factors for usability.

Determination of brightness, among other fluorescence properties is a standard procedure for characterisation of fluorescent proteins^20^. The method presented here does not rely on expression of a standard FP in the same cell, but a comparison to other fluorescent reporters is advisable. The gathered brightness data are specific to the used confocal imaging system. Yet, a comparison with published data using fluctuation spectroscopy techniques yields similar results on mEGFP versus mNeonGreen^30^ and mEGFP vs StayGold variants^31^. The characterisation of StayGold variants is needed as it is unclear how these variants perform in fair comparison. Evaluation using nanocages has yielded 3-fold higher brightness as compared to mEGFP with mSG outperforming StayGoldE138D and mBaoJin^42^. We observe similar trends but 7-fold higher brightness of mSG and mSG2 as compared to mEGFP. The reason for the discrepancies could be that the FCS analysis is evaluating brightness at the single molecule level whereas the nanocages rely on assembly of FPs and that all of them are folded and fluorescent. mSG and mSG2 show very similar brightness in our hands, which makes sense as they only differ in terminal extensions aiding folding and expression but not affecting the chromophore. Another study using FCS finds similar brightness between mSG and StayGoldE138D in cell culture and about two-fold higher brightness as compared to EGFP^31^. These inconsistencies might originate from the use of the fixed or limited range laser powers. We believe that by performing the FCS excitation scan, fitting the data and obtaining the usable brightness values will be more consistent and easier to compare, for example, by normalisation to maximum brightness. Notably, we compared the StayGold variants in mammalian cells after 24 h of transfection, studies in yeast^43^ and Drosophila^17^ report lower, similar brightness as compared to mNeonGreen, for instance.

Fitting FCS excitation scans with a saturation model for the brightness and a linear model for the transit time is convenient but represents a huge oversimplification. Photo-bleaching dynamics can be highly complex and the transit time against laser power data convolve saturation and photo bleaching effects along with other photophysical effects^44^. These phenomena are difficult to disentangle, especially in a biological specimen. However, as saturation in cpm in most cases outweighs changes to the transit time, we opted to keeping it simple and empirical. We are integrating over various effects and use the extracted brightness and laser power values with that in mind. Photophysical characteristics depend on the mode of illumination and the specific imaging instrument. Our FCS excitation scan pipeline has extensively been used on confocal setups with a specific set of lasers and optics. Future comparison to other setups, TIRF-FCS^45,46^ based, or SPIM-FCS^47^ based will allow us to generalise the method to other optical geometries.

The change of brightness with temperature is a well-known effect for RhB^48,49^, with an expected drop of 2.2% per centigrade increased temperature. The observed change in our hands from 28 °C to 37 °C is well within that prediction. Interestingly we also observe a decrease in brightness for the other dyes, AF488 and Atto655. This highlights the need to characterise probes under the conditions where the actual biological question is to be investigated. Our analysis pipeline uniquely allows the comparison of organic dyes, chemigenetic reporters, and fluorescent proteins on the same instrument and under the same conditions. This allows to put the measured brightness values directly in context.

The FCS analysis and fitting would ideally treat all data the same, i.e., fit with the same model. However, specifically for stagRFP and mScarlet as well as for the chemigenetic labels in the cells, we had to employ models with more fitting paraments: two triplet states (**Supplementary Figure S7**) or two diffusing components (**Supplementary Figure S10**), respectively. This might contribute to higher uncertainty of the diffusion coefficient and brightness values. It is advisable to keep the number of fitting parameters as low as possible. While these probes might be a reasonable choice for imaging, they should be used for fluctuation spectroscopy with caution.

This study focusses on brightness as determining factor for successful measurements with FCS^24,29^. However, also other parameters must be considered. For example, protein folding (maturation), oligomerisation, photo-stability, or photo-toxicity. All these parameters might vary between different organisms. For instance, FP performances have been compared in studies in C. elegans^18^, Drosophila^17^, yeast^16,43^, or mammalian cells^42^ with varying results probably due to different cellular milieus, expression levels and temperatures. Notably, the brightness values derived from FCS in our study correlate between zebrafish measured at 28 °C and HEK cells measured at 37 °C (**Figure 2k**). Thus, microscopy modalities and mode of measurement are important to consider. Lastly, some effects might not be immediately apparent but could be detrimental to the organism such as damage caused from reactive oxygen species generated by, for example, mCherry^50^. Careful characterizations need to be performed and multiple approaches integrated.

In conclusion, this study evaluates 10 FPs in zebrafish embryos, 14 in cell culture and 7 dyes in solution with 4 of them used for chemigenetic labelling. We cover a wide range of excitation and emission spectra as well as commonly used tags. However, new fluorophores are constantly being developed. We hope that availability of this workflow and analysis pipeline will encourage probe evaluation along with biological experiments. Overall, we present a pipeline that can be performed on any FCS capable microscope and permits comparisons of fluorescent probes in a quantitative fashion. Furthermore, we hope to start providing a good resource for researchers to pick a probe for their specific experiments.

## Acknowledgements

F.S. acknowledges support from the long-term postdoctoral fellowships support from EMBO (EMBO ALTF 849-2020) and HFSP (LT000404/2021-L) as well as from the Leverhulme Trust, UK (LIP-2021-017). All imaging has been performed at the Translational Imaging Center (TIC) at the University of Southern California. The authors thank Jason Junge and Arkadi Shwartz for technical assistance and microscope maintenance. The authors would like to thank Satyajit Mayor (University of Warwick) for careful proofreading and discussion of the manuscript as well as all the tool builders (FP engineers and chemical biologists) for developing the tested probes and making them available to the community.

## Author Contributions

F.S. and S.E.F. conceptualised the study. F.S. performed all the molecular work, experiments, and data analysis. L.A.T. provided guidance for the zebrafish work and molecular designs. F.S. wrote the manuscript under guidance by S.E.F. and L.A.T.

## Data Availability

All data are available upon reasonable request from the corresponding authors.

## Materials and Methods

### Zebrafish and ethics

Zebrafish lines were maintained according to standard practice^51^ and in accordance with the local regulations at the University of Southern California. The animal work is covered and approved by the IACUC protocol 12007 USC. Wildtype AB and TL lines were obtained from ZFIN. All experiments were performed on embryos from crossing theses lines. No lines have been raised.

### Plasmid design and construction

All plasmids have been designed and constructed using standard molecular biology techniques. The plasmids used in this study are summarised in **Table 1**. The pCS2+ dual purpose vector has been used for expression in mammalian cells (CMV promotor) as well as for in vitro transcription (SP6 promotor). The pCS2+ backbone was linearised with EcoRI-HF and SnaBI (NEB #R3101S and #R0130S). Fluorescent protein inserts have been PCR amplified or synthesized as geneblocks by IDT. An SV40 NLS signal (PKKKRKV) was placed at the N-terminus to localise the FP to the nucleus. All protein sequences have been checked against the sequence reported on FPbase.com^39^. PCRs were performed using La-Taq (Takara, #RR002A). Plasmid construction was performed using In-Fusion reaction (InFusion Snap Assembly #638947) according to the manufacturer’s protocol. Plasmid sequences were verified by Sanger sequencing and whole plasmid sequencing. Plasmids will be deposited in Addgene.

**Table 1:**
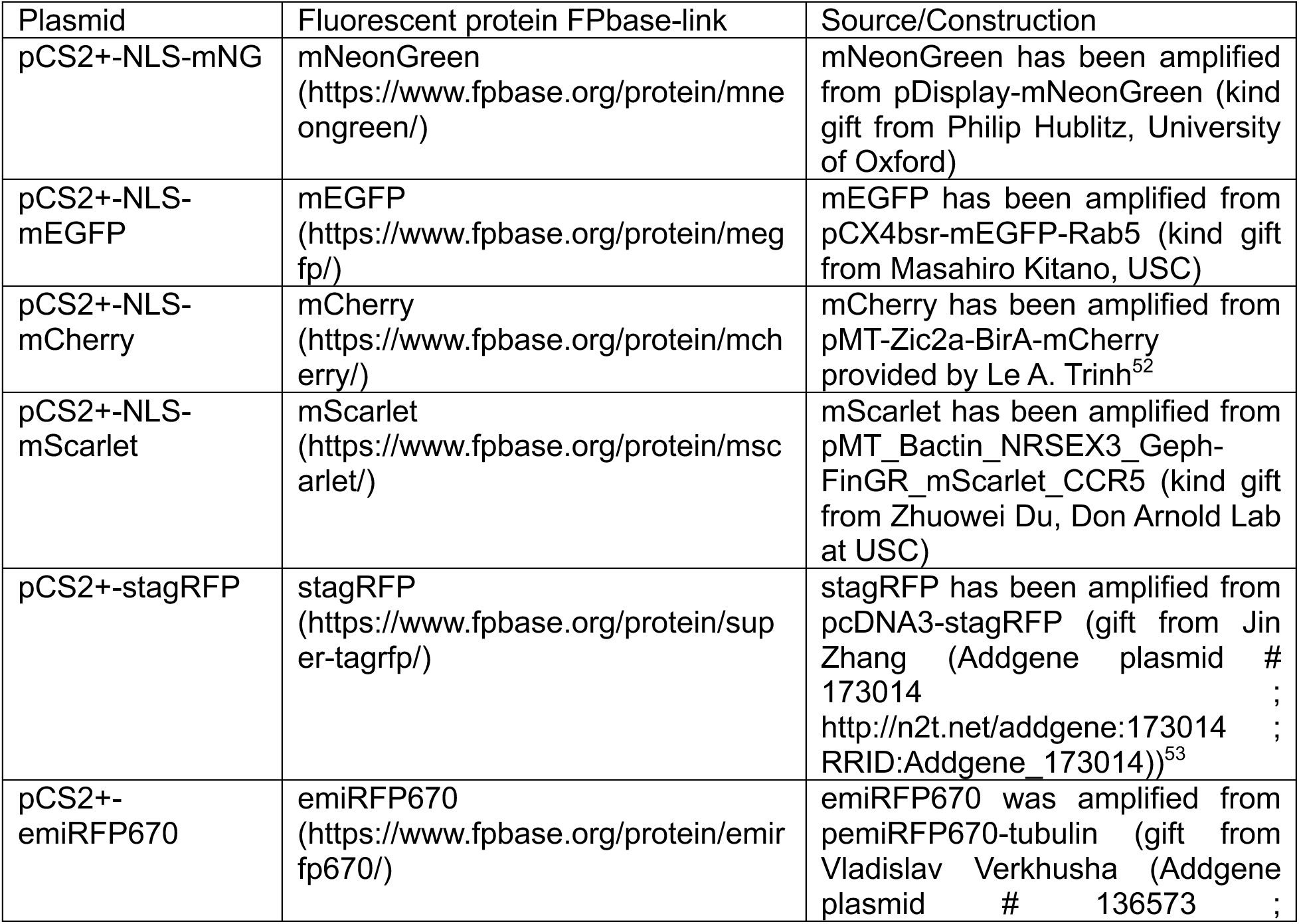
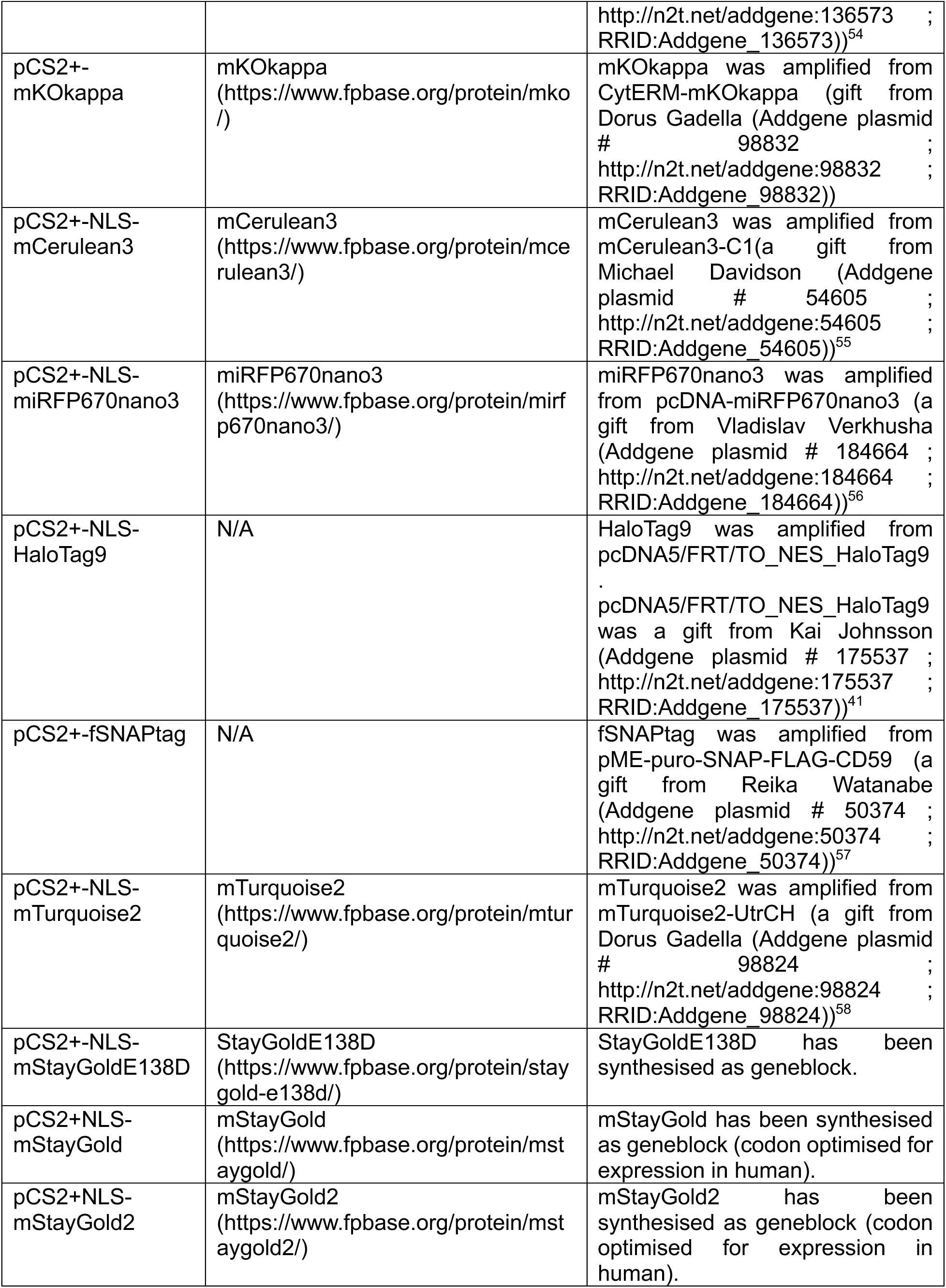
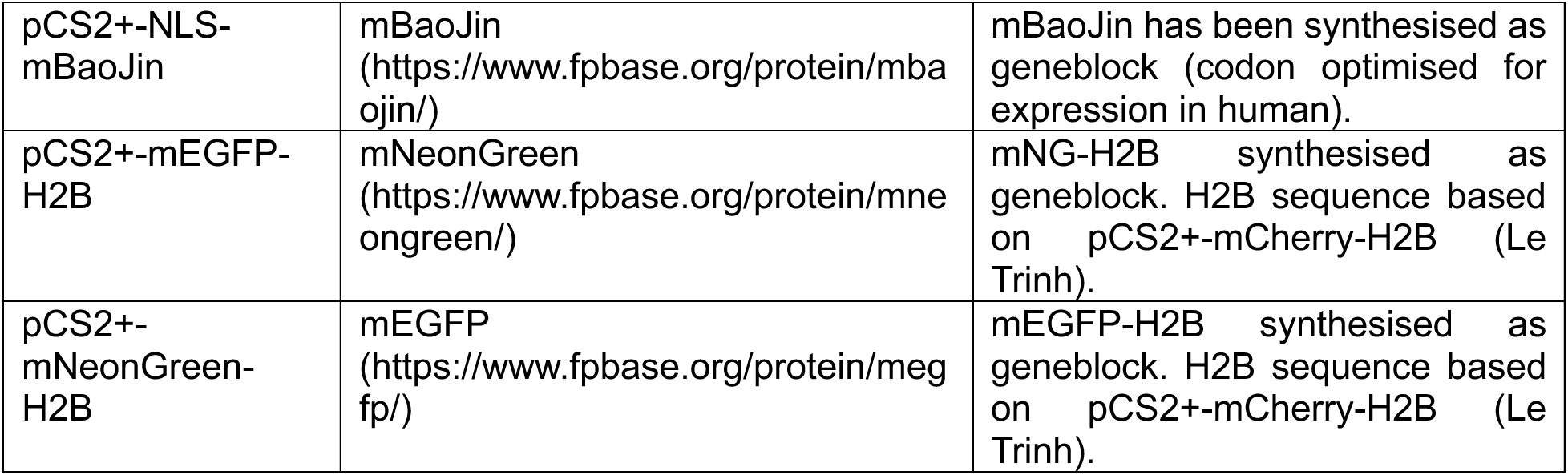
Plasmids used in this study.

### In vitro transcription and injection of mRNA

In vitro transcription was performed using the mMESSAGE mMACHINE™ SP6 Transcription Kit (Invitrogen, #AM1340) according to the manufacturer’s protocol. 5 µg of plasmid DNA were linearized with NotI-HF (NEB, #R0189). Note, pCS2+-NLS-mScarlet was linearized with with MfeI-HF (NEB, #R3589S) due to internal NotI restriction site. Linearised DNA was purified using QIAGEN PCR Clean-up kit (#28104) before in vitro transcription reaction. After transcription, mRNA was purified using QIAGEN RNAeasy kit (#74104). Samples were eluded with DNAse/RNAse free water (Ambion) and stored at 500 ng/µL in 2 µL aliquots at -80 °C. For microinjection, samples were diluted to 50 ng/µL (including 0.01% phenol red). 2.3 nL of mRNA solution were injected into single cell stage embryos using a nanoject II microinjector (Drummond Scientific) mounted on a micromanipulator. Injections were performed in the fish facility at 28 °C ambient temperature. Injected embryos and control embryos were kept at 28 °C till after gastrulation, dead embryos cleaned out, and petri dishes transferred to 23 °C to slow down development^59^.

### Cell Culture and Transfection

HEK293T cells were maintained according to standard practice at 37 °C and 5% CO_2_ in DMEM (Corning, #10-013-CV) supplemented with 10% fetal bovine serum (Thermo Fisher Scientific, #A5256701), 1% penicillin/streptomycin mix (Thermo Fisher Scientific, #15140122), and 1% L-glutamine (Sigma-Aldrich, #G7513). Cells were split twice a week and kept in T25 flasks. For imaging experiments, cells were seeded onto µ-Slide 8 well (ibidi, #80807) with #1.5 glass bottom. Plasmid transfection was performed with FuGENE 6 (Promega, #E2691) according to the manufacturer’s protocol. 50 ng of plasmid DNA were transfected per well (∼ 1cm^2^).

### Labelling of cells with HALO and SNAP ligands

HEK293T cells were maintained as described above. µ-Slides were coated with poly-D-Lysine (PDL, Gibco # A3890401) for 1 h and washed 5 times with PBS and full medium before seeding cells. Cells were labelled in full medium with HaloTag® Oregon Green® Ligand (Promega, G2802), Janelia Fluor® 646 HaloTag® Ligand (Promega, #HT1060), SNAP-Cell® 505-Star (NEB, #S9103S), or SNAP-Cell® 647-SiR (NEB, #S9102S). Cells were labelled at 0.5 µM final concentration in full medium for 1h and washed five times with full medium and phenol-red free Leibovitz 15, L15 (Gibco, #21083027), left to wash for 15 minutes and the procedure repeated. To increase washing efficiency and removal of unbound dye, cells were seeded on PDL coated channel slides (µ-Slide VI 0.5 Glass Bottom, ibidi # 80607).

### Sample preparation for microscopy

Dyes in solution (in vitro and calibration samples) were diluted in PBS (pH 7.4) to a concentration of 100 nM. HALO- and SNAP-ligands were diluted from frozen stock aliquots fresh on the day of experiments. Rhodamine B (Fisher Scientific # R21-100), Atto655-NHS (Sigma Aldrich # 76245-1MG-F), and AlexaFluor488-NHS (Thermo Fisher Scientific, # A20000) were prepared and stored at 4 °C. All measurements were performed in µ-Slide 8 well (ibidi, #80807) or µ-Slide 18 well (ibidi, # 81817) with #1.5 glass bottom. HEK293T cells were kept and transfected in full medium on glass bottom slides. Before imaging, full medium was replaced by prewarmed L15.

Zebrafish embryos were staged^59^. Embryos were manually screened for fluorescence on a stereoscope equipped with fluorescence lamp (Zeiss Axio Zoom). Low expression (i.e., dim) embryos were selected for FCS and higher expression for imaging. Embryos were dechorionated using fine forceps on a stereomicroscope and allowed to recover for a few hours. Imaging was performed on round glass bottom dishes (WillCo Wells #D5040P) with #1.5 glass thickness. 750 µL of 1% agarose (Invitrogen #16500-100) in 0.3xDanieau (17.4 mM NaCl, 0.21 mM KCl, 0.12 mM MgCl_2_, 0.18 Ca(NO_3_) 1.5 mM HEPE0S, pH 7.6) was used to coat the coverslip and a custom mold was pressed in, to create wells for the embryos. A solution of 0.01% Tricaine and 1% low-melt agarose (Invitrogen #6520-050) in 0.3xDanieau was added to the wells and embryos dropped right after. Embryos were oriented hindbrain towards coverslip using an eyelash pick. After gelation of the agarose the dish was topped up with 0.3xDanieu and closed to avoid evaporation on the microscope.

### Confocal microscopy and FCS

All imaging and FCS were performed on a Zeiss LSM 880 housed in the Translational Imaging Center and the instrument operated using the Zen Black software (Zeiss). The microscope was equipped with a Zeiss LD C-Apochromat 40x 1.1 NA water immersion objective. For excitation we utilised the following laser lines 458 nm, 488 nm, 561 nm, and 633 nm in combination with appropriate beam splitter 488/561/633 and 458/561 depending on the fluorophore’s excitation spectrum. For blue fluorophores emission was captured between 465-570 nm, for green fluorophores between 500-600 nm, for orange fluorophores 571-669 nm, and for red fluorophores 642-678 nm using the GaAsP hybrid detector (Channel S) running in photon counting mode. Laser powers were measured weekly using a power meter (Nova II, Ophir #7Z01550) as calibration for the FCS excitation scans and to confirm laser powers are linear within the measurement regime. In addition to fluorescence, brightfield from the transmission PMT was captured.

FCS data were captured using the FCS routine in Zen. The beam path was setup in the same way as in imaging. For every experiment the microscope was calibrated using a solution of fluorophore with known diffusion coefficient (Acridine Orange^60,61^, Alexa Fluor 488^62^, Rhodamine B^63^, Atto655^63^). Diffusion coefficients were recalculated for measurement temperature where literature values were not available according to^63,64^. The same dye solution has been used to align the pinhole and to adjust the correction collar as described in detail in^65^. The microscope was set to 28 or 37 °C and along with the calibration solution to equilibrate for 1 h prior to any measurements. Intensity traces were recorded and hardware correlated. Data were saved as .fcs files. These correlated FCS data were analysed using the FoCuS_point software package (version 1_16_208)^66^. FCS data were fitted in the range from 1 µs to 1 s to a function of the general form:

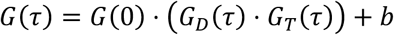

Where G is the correlation value at give lag time, *τ*. The correlation amplitude G(0) is inversely proportional to the average number of molecules in focus. *G_D_*(*τ*) describes the diffusive processes in the sample and *G_T_*(*τ*) describes the photophysical processes, e.g., triplet state transitions. b is an offset.

All curves obtained were fitted to a 3D diffusion model:

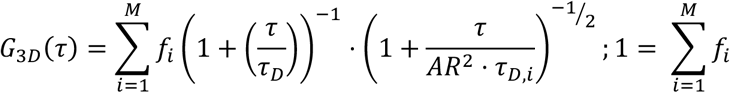

Where *τ_D_* is the transit time, average time for a fluorescent particle to transverse the observation volume, *f_i_* is the fraction of component i (in a multicomponent fit), and *AR* is the aspect ratio between long and short axis of the observation volume. All data on dyes in solution, fluorescent proteins in cells or zebrafish embryos were fitted with a single component diffusion model (i=1). Data on HALO and SNAP ligands in cells were fitted with a two-component model (i=2). The dark state kinetics were described as follows:

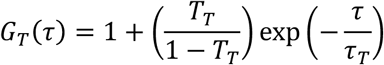

Here *τ_T_* is the triplet time and *T_T_* the triplet fraction. Triplet times were typically 5 µs for dyes and 40 µs for fluorescent proteins^24^. Note for data on stagRFP and mScarlet, we included a second long-lived dark state with a decay time of around 150 µs.

The calibration data were used to recalculate the beam waist radius, *ω*, according to

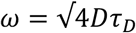

Where D is the diffusion coefficient reported in the literature (as described above). Fits were exported from the FoCuS_point software package and stored in Excel sheets for further downstream analysis.

### FCS excitation scans

To obtain one FCS excitation scan for analysis a suitable area within the hindbrain at a depth of 50-60 µm from first cell layer was chosen. Measurements from multiple locations were pooled. Specifically for measurements on embryos, 3 repeats of curves at three different locations within three different nuclei were acquired at varying laser powers. Thus, every embryo measurement consisted of 9 FCS curves per laser power. The data were collated per embryo. For all FPs, three different biological repeats, ie three independent sets of crossings and injection, were obtained with at least 5 embryos each. The data pooled over multiple embryos (ie one biological repeat) was analysed. For data obtained in HEK cells, three different locations per nucleus were measured for every laser power. At least 8 nuclei were measured per independent transfection (ie biological repeat). Pooling the data helped establish robustness of the measurements against inherent biological noise.

FCS excitation scan data consisted of cpm (in kHz) and transit time (in ms) versus incident laser power (in µW). These data were analysed with a custom Python script available on GitHub (https://github.com/Faldalf/FCS-Exc-Scan). In brief, the brightness was fitted with a saturation model of the form:

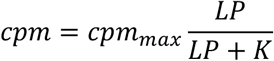

Where *cpm_max_* is the maximum brightness, *LP* is the laser power, and *K* is the inflection point of the saturation curve. The changes to transit time, either an increase due to optical saturation or a decrease due to photobleaching have been fitted to a linear model. Main output of the fitting routine is the useable brightness *cpm*_*use* and the useable laser power *LP_use_*. This point is defined as either 10% of the inflection point K in the saturation curve of the cpm against laser power or a 20% change to the initial value of the transit time. In our hands, the saturation of cpm typically determines the useable parameters. Automated fitting and data handling was facilitated by use of scipy^67^ and pandas^68^.

### z-scanned FCS

FCS data at fixed laser power but varying depth were acquired in a similar fashion as described previously. The 0-depth position was defined as the first layer of cells. Using the Zeiss LSM880 stage z-controller three repetitions of FCS data were acquired for every nucleus at incrementally increasing depth, every few µm. At the end of the experiment, the 0-depth point was verified by imaging the same area again. For every FP at least 5 embryos were measured and the resulting FCS data vs normalised depth concatenated and processed using a sliding window approach. Code available on GitHub (https://github.com/Faldalf/FCS-Exc-Scan).

### Image processing

Imaging data from the Zeiss880 confocal were saved as lsm5, opened and processed using ImageJ/FIJI^69^. Images were only minimally processed by adjusting brightness/contrast, cropping if larger field of view acquired or scale bars added.

### Statistics

Statistical comparisons were performed using GraphPad Prism. Comparisons were always performed on pooled data as described above. Useable brightness and laser power were compared between groups of FPs with the same excitation wavelength. I.e., green FPs were excited at 488 nm and compared to each other. For pairwise comparisons a student’s t-test was employed. For multiple comparisons a one-way ANOVA with Tukey’s correction was used. Statistical comparison of values across different groups (e.g., useable brightness of red FPs versus green FPs) is provided in Supplementary Table T1-T4 using one-way ANOVA with Tukey’s correction.

