## Supplementary Information for "Quantitative comparison of fluorescent reporters by FCS excitation scan"

### Title

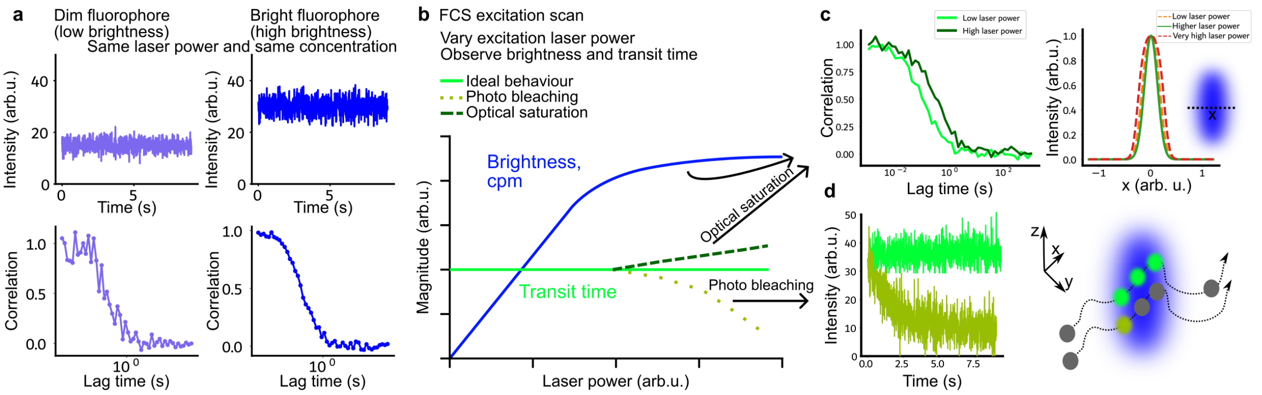


### Supplementary Figure S1: Schematics of FCS excitation properties

**a** Cartoon of intensity trace (top) and autocorrelation curve (bottom) of a fluorophore with low brightness (left) or high brightness (right). At the same concentration and the same excitation laser power, a bright fluorophore will yield more photons per unit time, higher intensity. This also improves the signal-to-noise ratio (scatter) of the autocorrelation curve.

**b** Cartoon of FCS excitation scan plot. FCS fitting parameters, brightness and transit time, are plotted against excitation laser power. In ideal case transit time does not change with laser power. Photobleaching or optical saturation can cause deviations.

**c** Optical saturation leads to an effective increase in the observation volume because the fluorescence response becomes non-linear with excitation intensity (right). Since saturation occurs first at the center of the focus, the effective fluorescence profile broadens. In the autocorrelation curves (left), this manifests as a shift to longer transit times.

**d** Photobleaching leads to irreversible loss of fluorescence from a fluorophore, resulting in a gradual decrease in signal intensity. If bleaching occurs on a timescale comparable to molecular diffusion through the observation volume, molecules may bleach before diffusing out. This introduces an additional decay process in the autocorrelation function and can lead to an artificially shortened apparent transit time in FCS analysis.

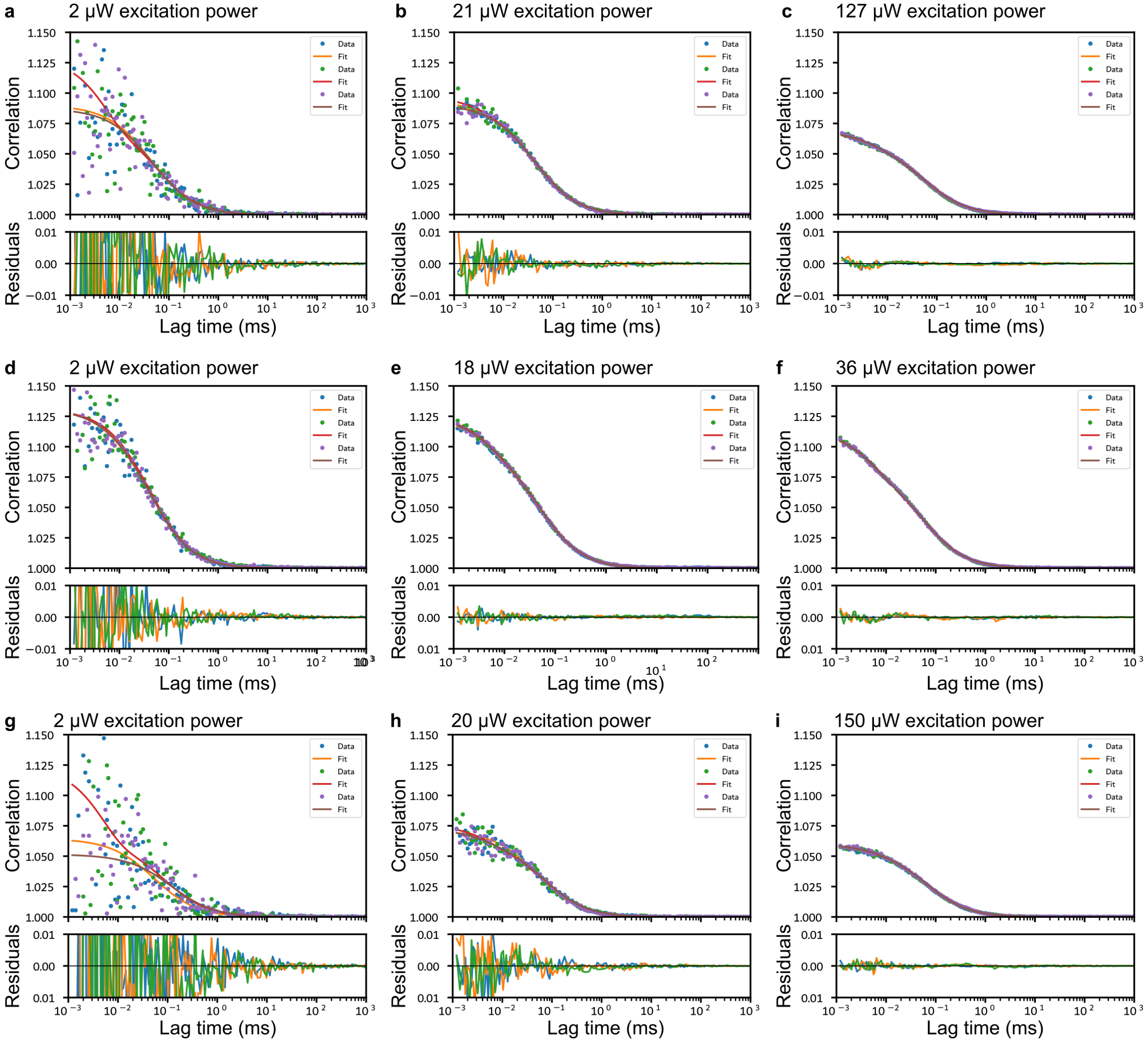


**Supplementary Figure S2: Examples of FCS raw data and fits for Rhodamine B, Alexa Fluor 488, and Atto655**

FCS data and fits with residuals (bottom panels) for

**a-c** Rhodamine B (100 nM in water) at 2, 21,127 µW 561 nm excitation laser power

**d-f** AlexaFluor 488 (150 nM in water) at 2, 18, 36 µW 488 nm excitation laser power

**g-i** Atto655 (150 nM in water) at 2, 20, 150 µW 633 nm excitation laser power

Shown are three repetition per dataset with three fits and respective fitting residuals.

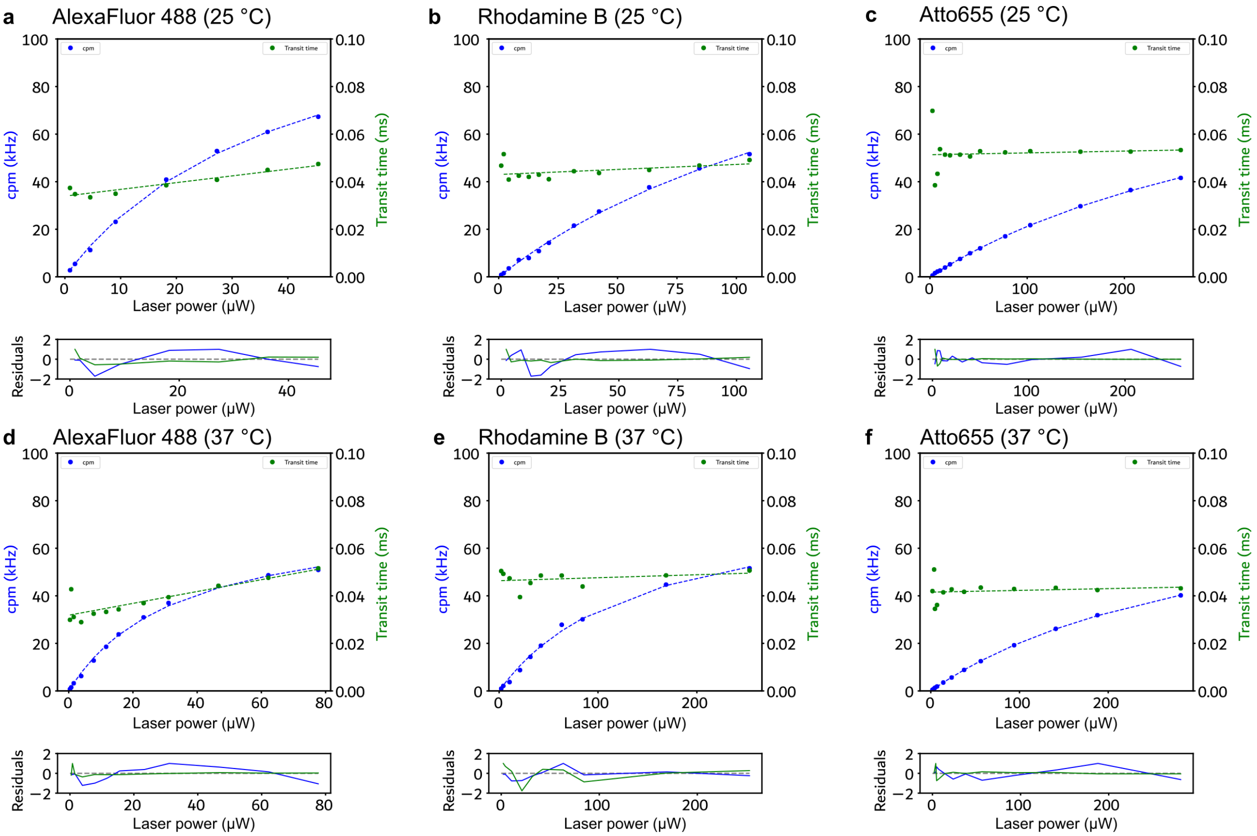


**Supplementary Figure S3: Examples of FCS excitation scan fits for Alexa Fluor 488, Rhodamine B and Atto655 at room temperature and at 37 °C**

Increase in brightness (blue) and change to transit time (green) with fitted saturation model for brightness and linear model for transit time shown as dashed lines. Bottom panels show fitting residuals.

**a** AlexaFluor 488 at 25 °C (150 nM in water)
**b** Rhodamine B at 25 °C (100 nM in water)
**c** Atto655 at 25 °C (150 nM in water)
**d** AlexaFluor 488 at 37 °C (150 nM in water)
**e** Rhodamine B at 37 °C (100 nM in water)
**f** Atto655 at 37 °C (150 nM in water)

Every dot represents the average over 3 FCS acquisitions.

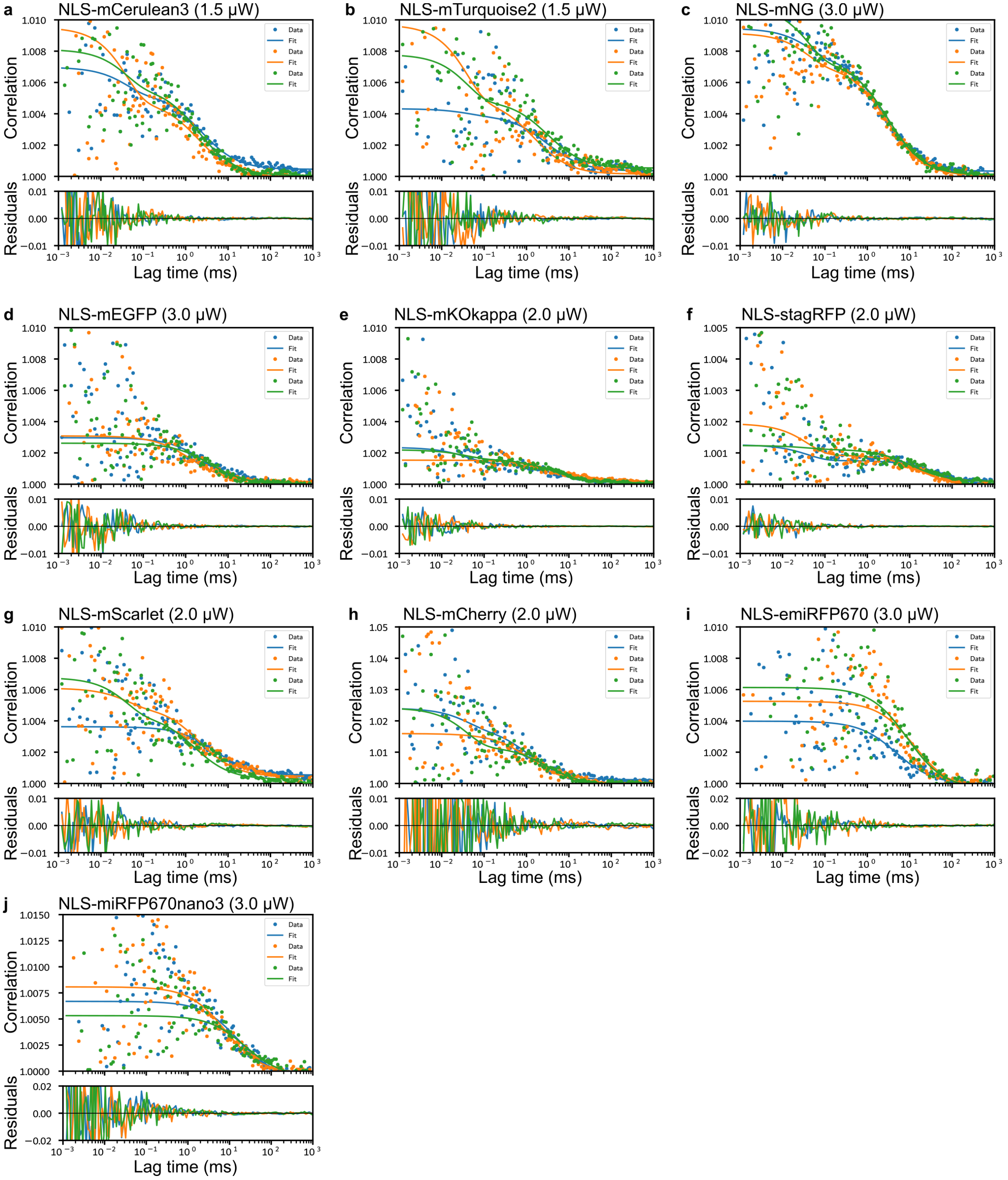


**Supplementary Figure S4: Examples of FCS raw data and fits for fluorescent proteins in zebrafish hindbrain at ~50 μm depth at 28 °C**

Shown are examples of raw correlation data (circles) and fitted curves (solid lines) for different acquisitions at indicated excitation power. Bottom panels show residuals.

**a** NLS-mCerulean3

**b** NLS-mTurquoise2

**c** NLS-mNeonGreen
**d** NLS-mEGFP
**e** NLS-mKOkappa
**f** NLS-stagRFP
**g** NLS-mScarlet
**h** NLS-mCherry
**i** NLS-emiRFP670
**j** NLS-miRFP670nano3

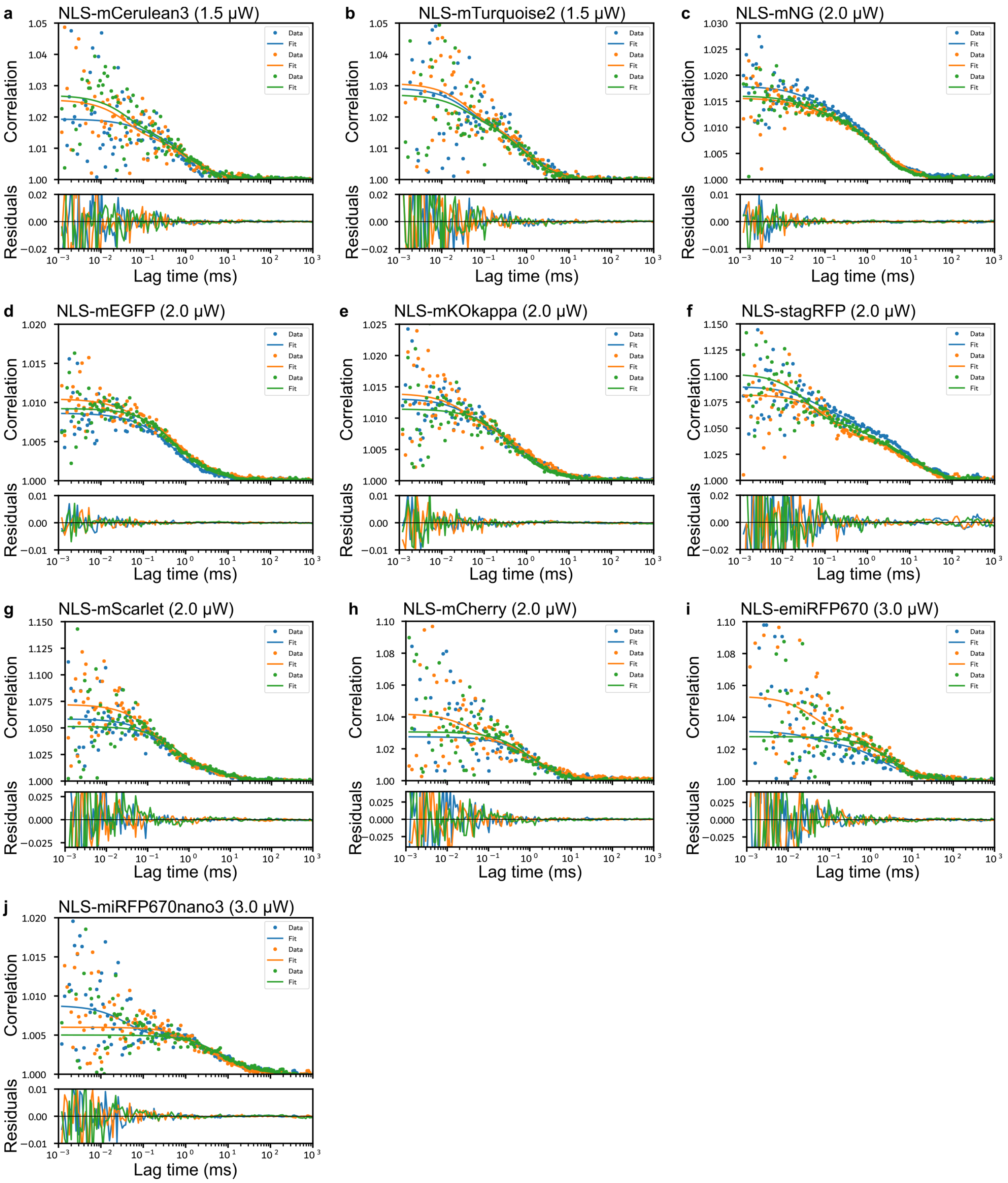


**Supplementary Figure S5: Examples of FCS raw data and fits for fluorescent proteins in HEK 293T cells at 28 °C**

Shown are examples of raw correlation data (circles) and fitted curves (solid lines) for different acquisitions at indicated excitation power. Bottom panels show residuals.

**a** NLS-mCerulean3
**b** NLS-mTurquoise2
**c** NLS-mNeonGreen
**d** NLS-mEGFP
**e** NLS-mKOkappa
**f** NLS-stagRFP
**g** NLS-mScarlet
**h** NLS-mCherry
**i** NLS-emiRFP670
**j** NLS-miRFP670nano3

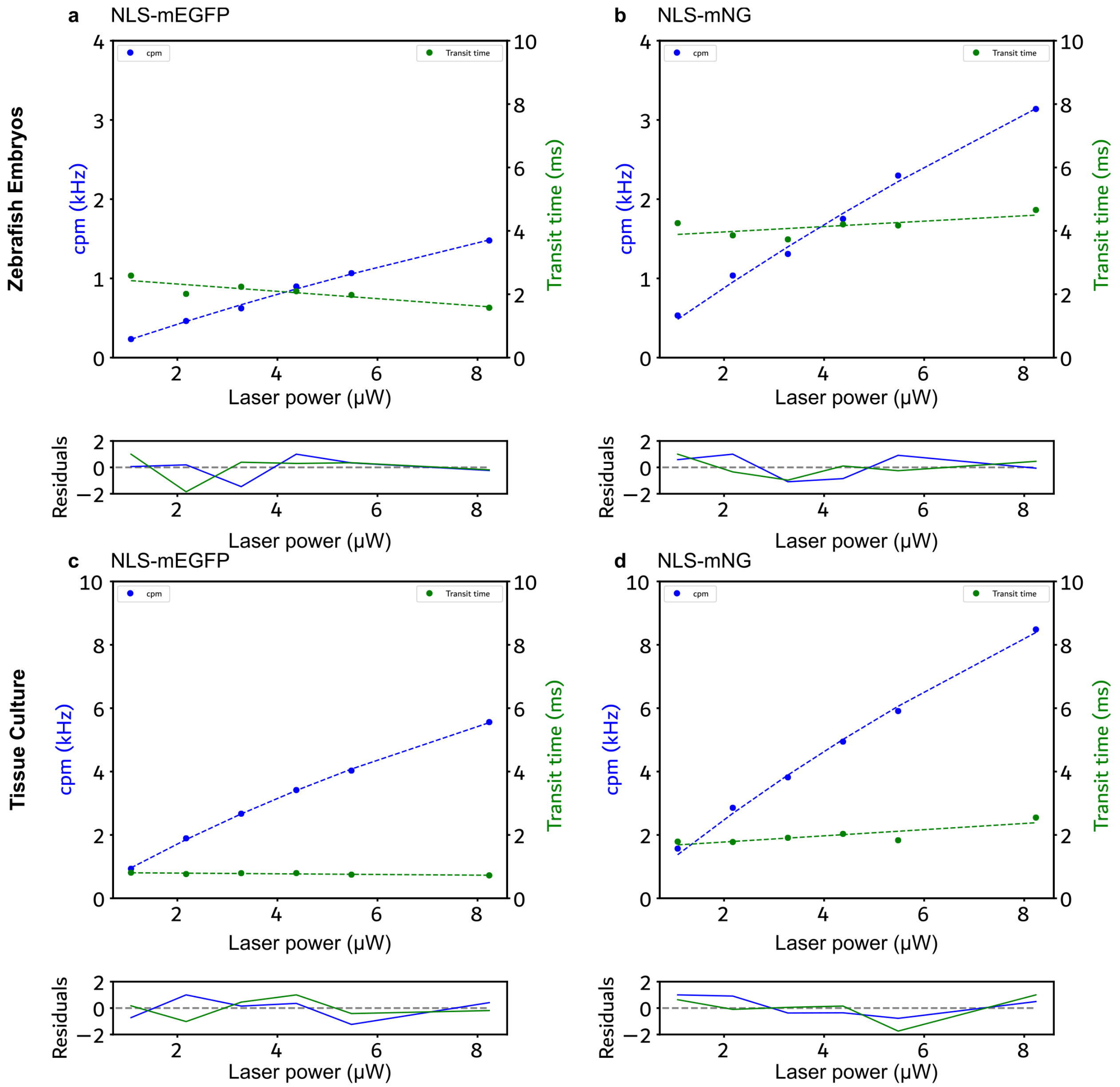


**Supplementary Figure S6: Examples of fitted FCS excitation scans for mEGFP and mNG acquired in zebrafish embryos and tissue culture.**

Shown are examples of cpm (blue) and transit time (green) against excitation laser power (488 nm) and fits (dashed lines). Bottom panels show residuals.

**a** NLS-mEGFP in zebrafish hindbrain at 28 °C
**b** NLS-mNG in zebrafish hindbrain at 28 °C
**c** NLS-mEGFP in tissue culture (HEK293T cells) at 37 °C
**d** NLS-mNG in tissue culture (HEK293T cells) at 37 °C

Every dot represents the average over multiple FCS acquisitions in 3 nuclei of at least 8 embryos or cells each.

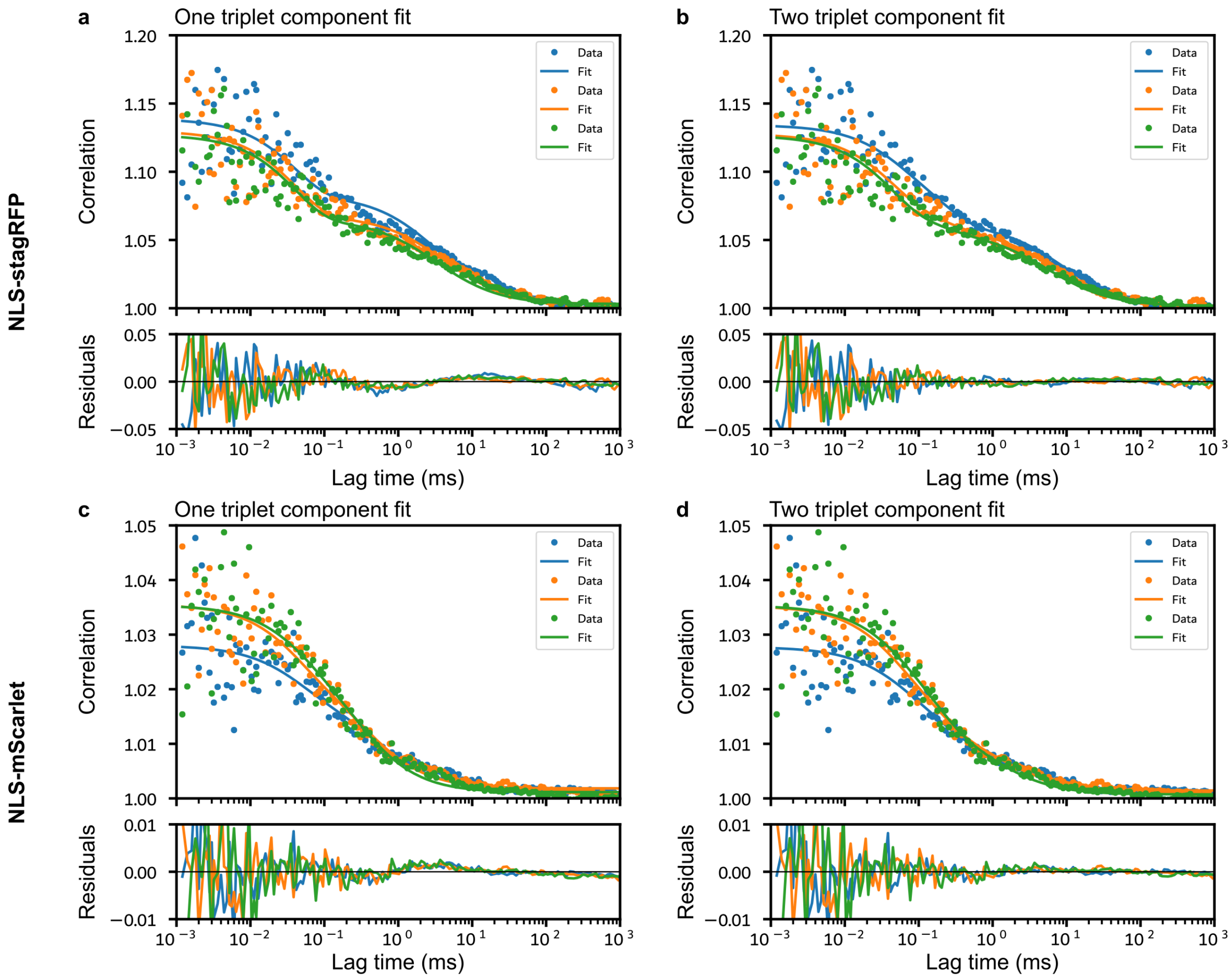


**Supplementary Figure S7: stagRFP and mScarlet show complex dark state kinetics.**

Shown are raw FCS data and fits to a model including one or two triplet state components (see Materials and Methods for details). Data are from transfected HEK296T cells acquired at 37 °C and at 4 μW excitation power (561 nm). Dots are raw FCS data and solid lines are fits. Bottom panels show fitting residuals.

**a** NLS-stagRFP data fitted to a model including one dark state component
**b** NLS-stagRFP data fitted to a model including two dark state components
**c** NLS-mScarlet data fitted to a model including one dark state component
**d** NLS-mScarlet data fitted to a model including two dark state components

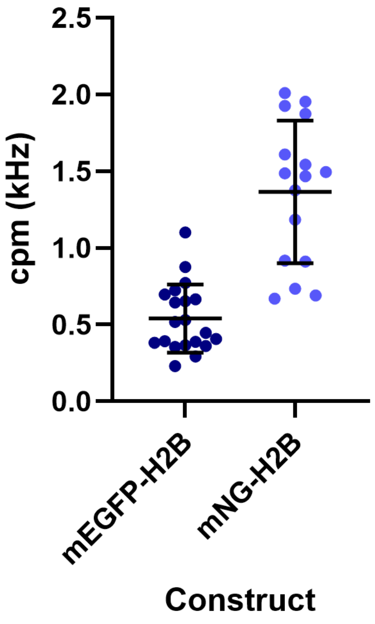


**Supplementary Figure S8: Comparison of cpm (brightness) of H2B tagged construct in zebrafish hindbrain**

Every dot represents one nucleus and averaging over 3-5 FCS curves. Data were acquired on 6 individual zebrafish embryos. Horizontal line represents mean, bars are standard deviation.

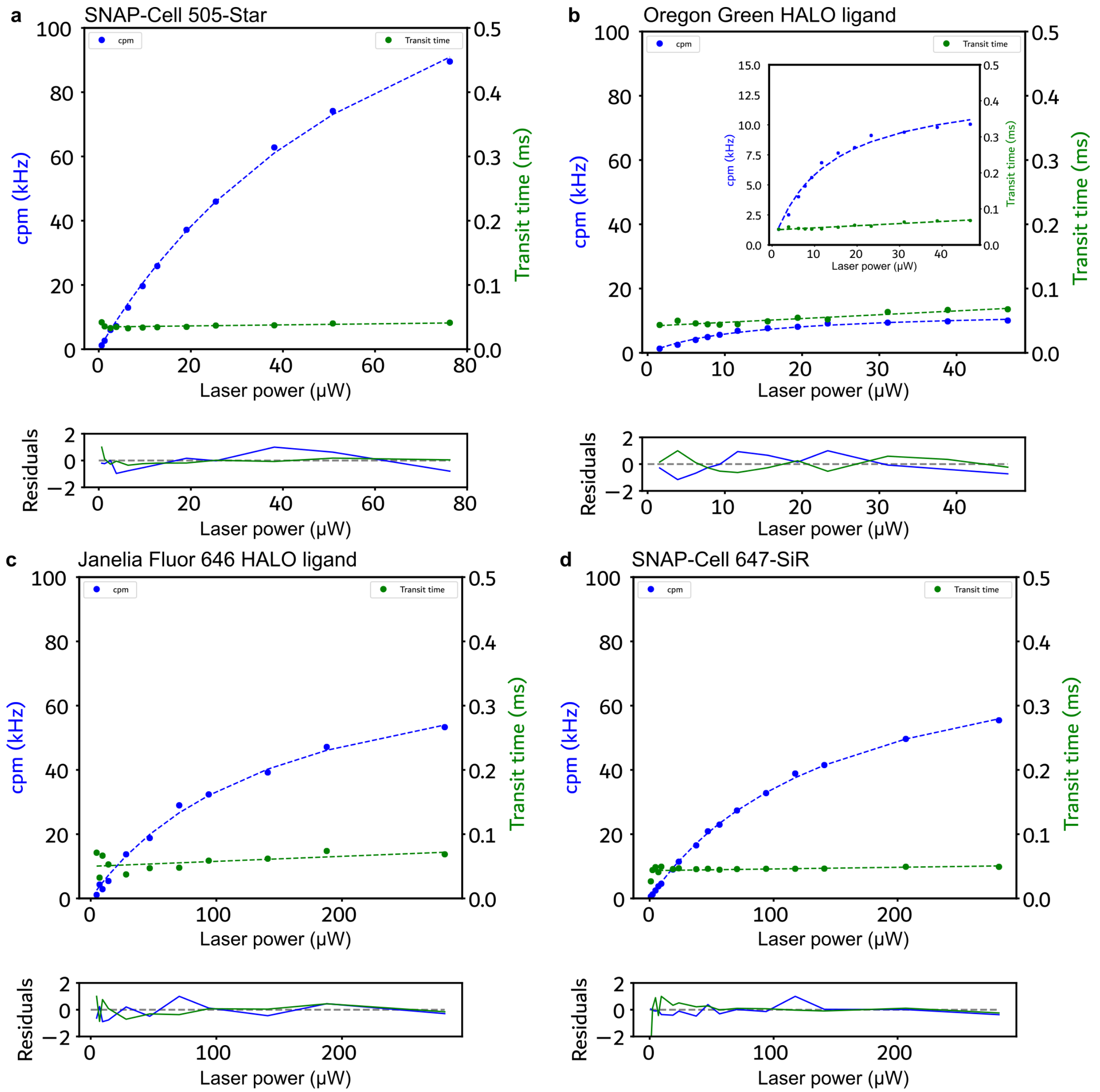


**Supplementary Figure S9: Examples of FCS excitation scans of free HALO and SNAP ligands in solution at 37 °C**

Shown are examples of cpm (blue) and transit time (green) against excitation laser power (488 nm for green dyes and 633 nm for far red dyes) and fits (dashed lines). Bottom panels show residuals.

**a** SNAP-Cell 505-Star
**b** Oregon Green HALO ligand (inset shows zoom in of the same data)
**c** Janelia Fluor 646 HALO ligand
**d** SNAP-Cell 647SiR

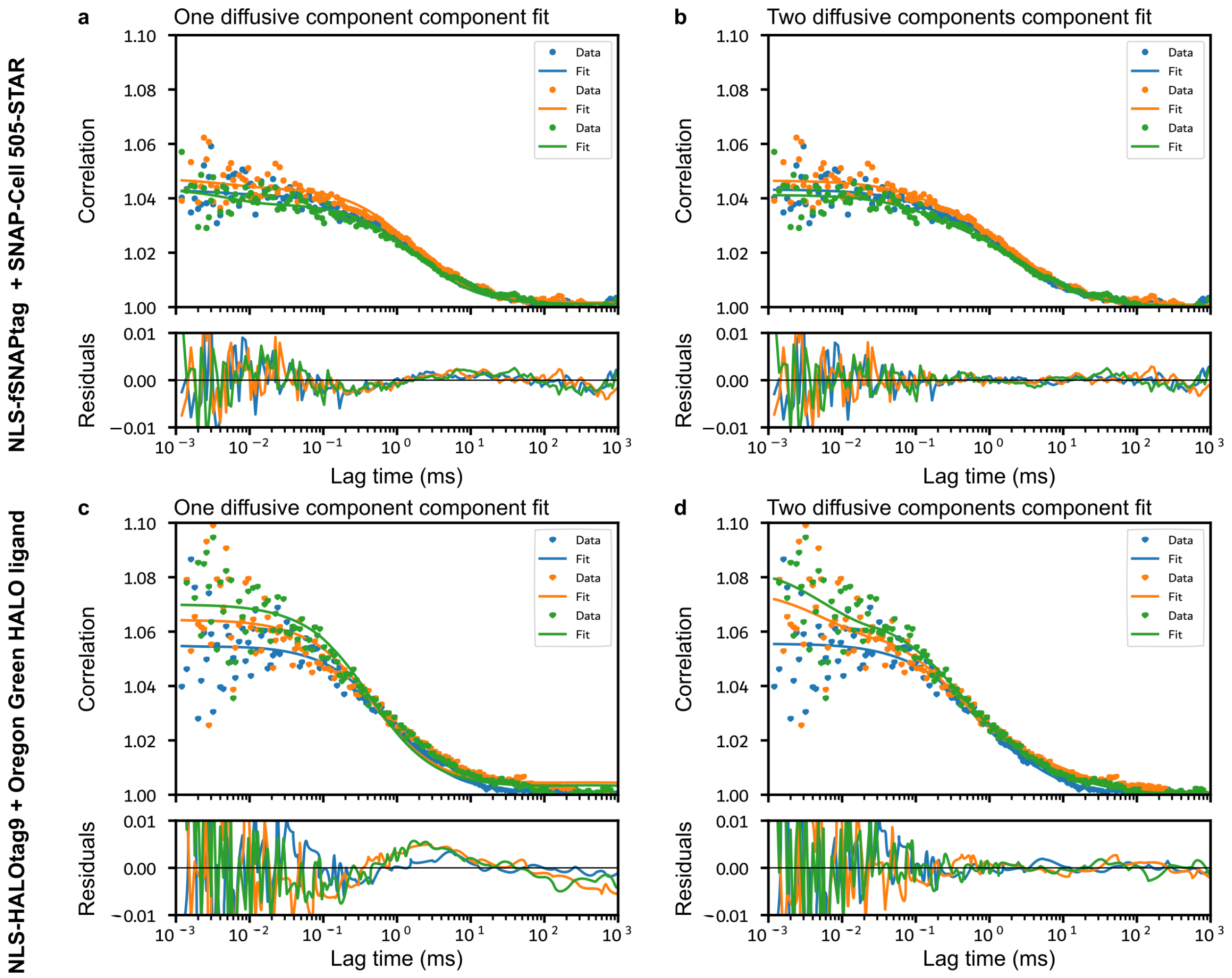


**Supplementary Figure S10: NLS-fSNAPtag and NLS-HALOtag9 require two component fit.**

Shown are raw FCS data and fits to a model including one or two diffusive components (see Materials and Methods for details). Data are from transfected HEK296T cells acquired at 37 °C and at 5 µW excitation power (488 nm). Dots are raw FCS data and solid lines are fits. Bottom panels show fitting residuals.

**a** NLS-fSNAPtag labelled with SNAP-Cell 505-STAR data fitted to a model including one diffusive component
**b** NLS-fSNAPtag labelled with SNAP-Cell 505-STAR data fitted to a model including two diffusive components
**c** NLS-HALOtag9 labelled with Oregon Green HALO ligand data fitted to a model including one diffusive component
**d** NLS-HALOtag9 labelled with Oregon Green HALO ligand data fitted to a model including two diffusive components

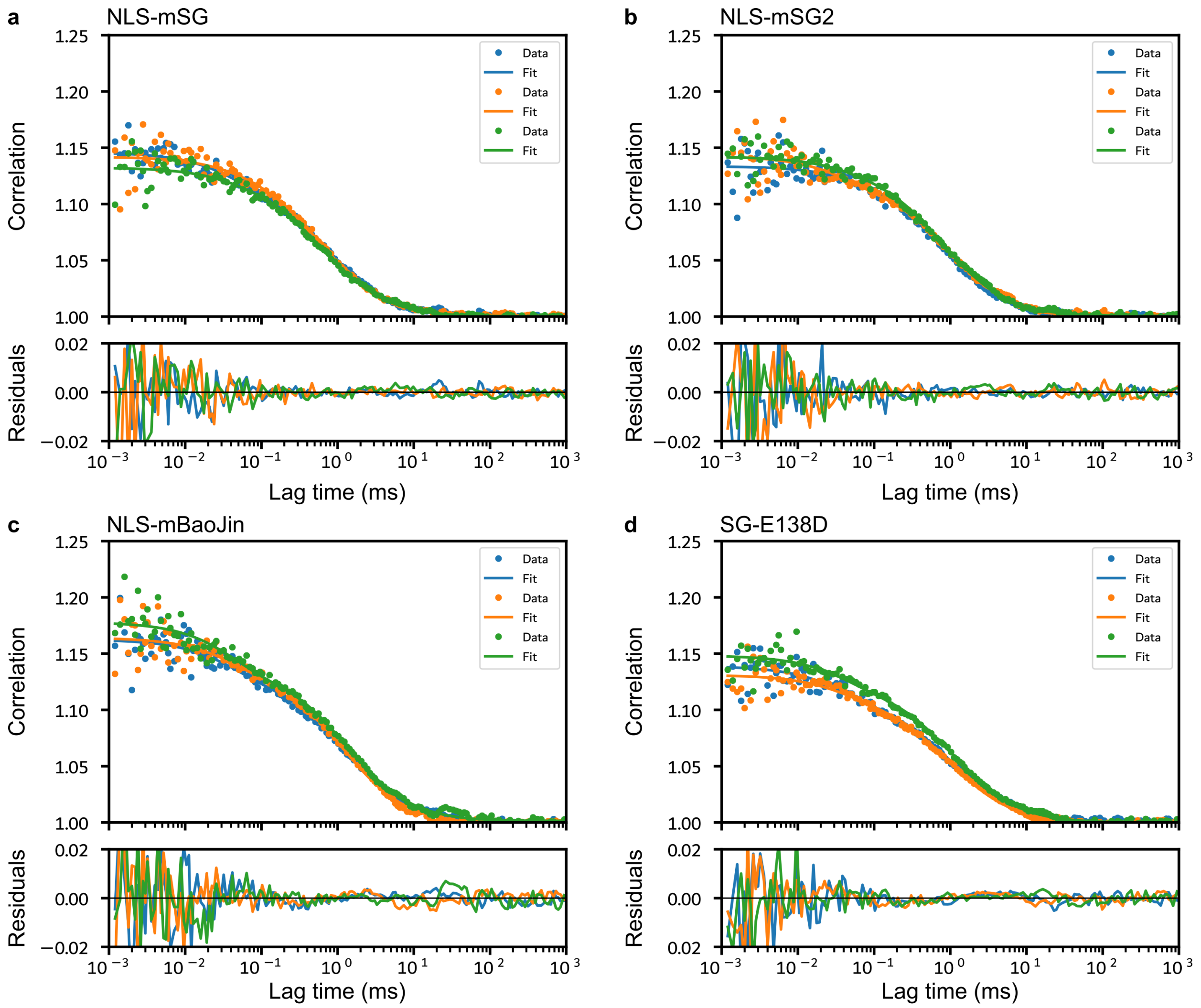


**Supplementary Figure S11: Examples of FCS curves and fits for monomeric StayGold variants.**

Shown are raw FCS data and fits (see Materials and Methods for details). Data are from transfected HEK296T cells acquired at 37 °C and at 5 µW excitation power (488 nm). Dots are raw FCS data and solid lines are fits. Bottom panels show fitting residuals.

**a** NLS-mSG
**b** NLS-mSG2
**c** NLS-mBaoJin
**d** NLS-SG-E138D

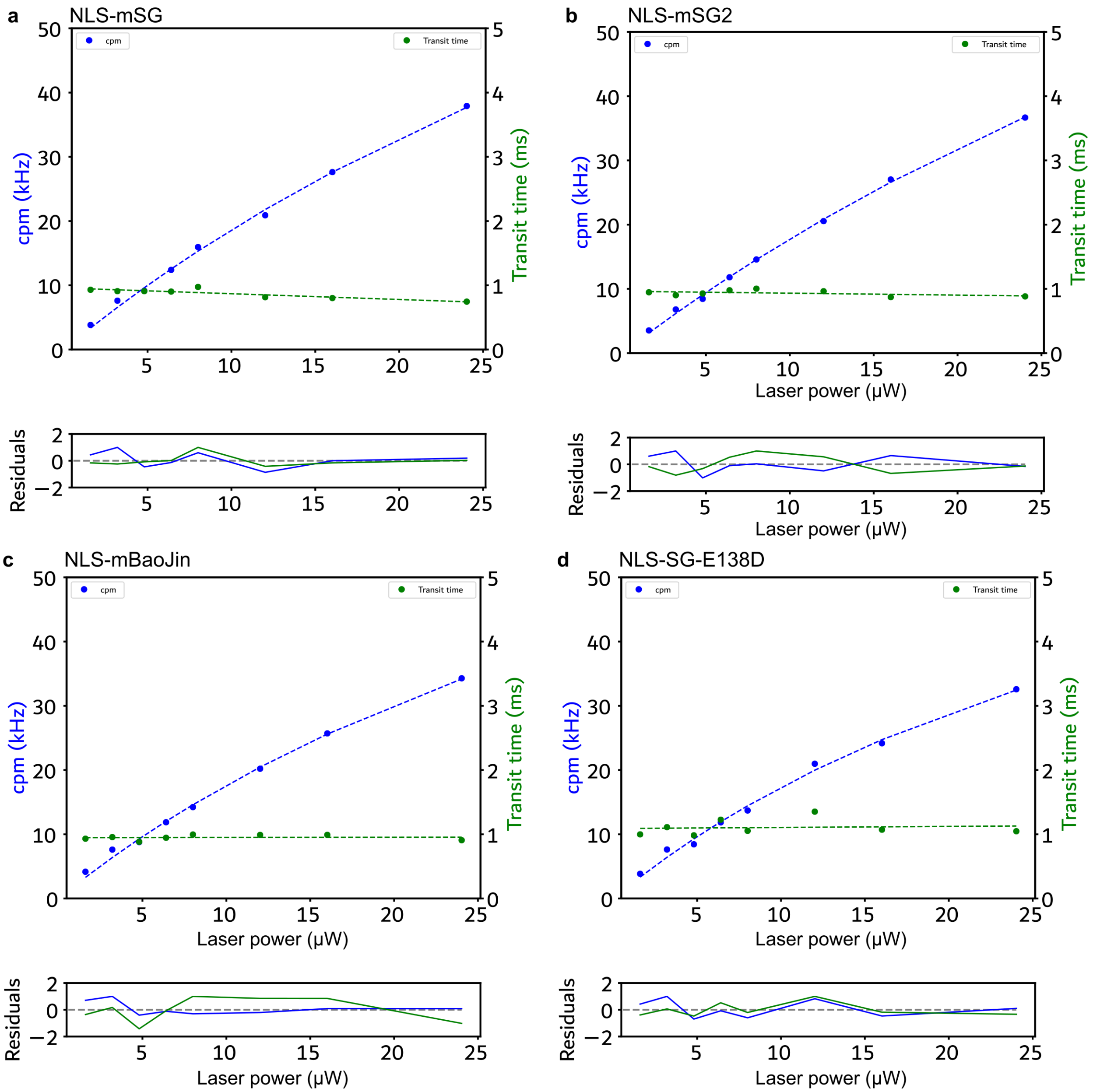


**Supplementary Figure S12: Examples of FCS excitation scans of the monomeric StayGold variants expressed with NLS-tags in HEK239T cells measured at 37 °C**

Shown are examples of cpm (blue) and transit time (green) against excitation laser power (488 nm) and fits (dashed lines). Bottom panels show residuals.

**a** NLS-mSG
**b** NLS-mSG2
**c** NLS-mBaoJin
**d** NLS-SG-E138D

**Supplementary Table T1**: Ajusted p-values from one-way ANOVA with Tukey's correction for multiple comparisons for useable brightness of FPs in zebrafish embryos. Plot in Figure 2g.

| | NLS-mCerulean3 | NLS-mTurquoise2 | NLS-mEGFP | NLS-mNG | NLS-mKOkappa | NLS-stagRFP | NLS-mScarlet | NLS-mCherry | NLS-emiRFP670 | NLS-miRFP670nano3 |
|---|---|---|---|---|---|---|---|---|---|---|
| NLS-mCerulean3 | | | | | | | | | | |
| NLS-mTurquoise2 | 0.99 | | | | | | | | | |
| NLS-mEGFP | 0.0002 | <0.0001 | | | | | | | | |
| NLS-mNG | <0.0001 | <0.0001 | <0.0001 | | | | | | | |
| NLS-mKOkappa | >0.9999 | 0.9689 | 0.0006 | <0.0001 | | | | | | |
| NLS-stagRFP | 0.9997 | >0.9999 | <0.0001 | <0.0001 | 0.9868 | | | | | |
| NLS-mScarlet | 0.9986 | >0.9999 | <0.0001 | <0.0001 | 0.9685 | >0.9999 | | | | |
| NLS-mCherry | >0.9999 | 0.9921 | 0.0001 | <0.0001 | >0.9999 | 0.9979 | 0.9920 | | | |
| NLS-emiRFP670 | >0.9999 | 0.9448 | 0.0008 | <0.0001 | >0.9999 | 0.9730 | 0.9441 | >0.9999 | | |
| NLS-miRFP670nano3 | 0.1489 | 0.0334 | 0.1425 | <0.0001 | 0.3026 | 0.0453 | 0.0332 | 0.1344 | 0.3611 | |

**Supplementary Table T2**: Adjusted p-values from one-way ANOVA with Tukey's correction for multiple comparisons for useable brightness of FPs in tissue culture. Plot in Figure 2h.

| | NLS-mCerulean3 | NLS-mTurquoise2 | NLS-mEGFP | NLS-mNG | NLS-mKOkappa | NLS-stagRFP | NLS-mScarlet | NLS-mCherry | NLS-emiRFP670 | NLS-miRFP670nano3 |
|---|---|---|---|---|---|---|---|---|---|---|
| NLS-mCerulean3 | | | | | | | | | | |
| NLS-mTurquoise2 | 0.0004 | | | | | | | | | |
| NLS-mEGFP | 0.9999 | 0.0005 | | | | | | | | |
| NLS-mNG | <0.0001 | <0.0001 | <0.0001 | | | | | | | |
| NLS-mKOkappa | <0.0001 | 0.8950 | <0.0001 | <0.0001 | | | | | | |
| NLS-stagRFP | <0.0001 | 0.0300 | <0.0001 | <0.0001 | 0.4330 | | | | | |
| NLS-mScarlet | <0.0001 | 0.0562 | <0.0001 | <0.0001 | 0.6124 | >0.9999 | | | | |
| NLS-mCherry | <0.0001 | 0.4119 | <0.0001 | <0.0001 | 0.9984 | 0.7959 | 0.9265 | | | |
| NLS-emiRFP670 | <0.0001 | 0.4458 | <0.0001 | <0.0001 | 0.9976 | 0.8867 | 0.9676 | >0.9999 | | |
| NLS-miRFP670nano3 | 0.0418 | 0.5459 | 0.0561 | <0.0001 | 0.0449 | 0.0002 | 0.0005 | 0.0047 | 0.0075 | |

**Supplementary Table T3**: Adjusted p-values from one-way ANOVA with Tukey's correction for multiple comparisons for useable laser power of FPs in zebrafish embryos. Plot in Figure 2i.

| | NLS-mCerulean3 | NLS-mTurquoise2 | NLS-mEGFP | NLS-mNG | NLS-mKOkappa | NLS-stagRFP | NLS-mScarlet | NLS-mCherry | NLS-emiRFP670 | NLS-miRFP670nano3 |
|---|---|---|---|---|---|---|---|---|---|---|
| NLS-mCerulean3 | | | | | | | | | | |
| NLS-mTurquoise2 | 0.999 | | | | | | | | | |
| NLS-mEGFP | 0.0211 | 0.0041 | | | | | | | | |
| NLS-mNG | 0.0218 | 0.0042 | 0.999 | | | | | | | |
| NLS-mKOkappa | 0.999 | 0.9999 | 0.0094 | 0.0097 | | | | | | |
| NLS-stagRFP | 0.9875 | 0.9999 | 0.0022 | 0.0023 | 0.9996 | | | | | |
| NLS-mScarlet | 0.9869 | 0.9999 | 0.0021 | 0.0022 | 0.9995 | 0.9999 | | | | |
| NLS-mCherry | 0.9954 | 0.7955 | 0.0759 | 0.0782 | 0.9478 | 0.6249 | 0.6195 | | | |
| NLS-emiRFP670 | 0.0819 | 0.0173 | 0.999 | 0.9996 | 0.0385 | 0.0094 | 0.0092 | 0.2678 | | |
| NLS-miRFP670nano3 | <0.0001 | <0.0001 | 0.0093 | 0.0090 | <0.0001 | 0.0001 | <0.0001 | <0.0001 | 0.0021 | |

**Supplementary Table T4**: Adjusted p-values from one-way ANOVA with Tukey's correction for multiple comparisons for useable laser power of FPs in tissue culture. Plot in Figure 2j.

| | NLS-mCerulean3 | NLS-mTurquoise2 | NLS-mEGFP | NLS-mNG | NLS-mKOkappa | NLS-stagRFP | NLS-mScarlet | NLS-mCherry | NLS-emiRFP670 | NLS-miRFP670nano3 |
|---|---|---|---|---|---|---|---|---|---|---|
| NLS-mCerulean3 | | | | | | | | | | |
| NLS-mTurquoise2 | 0.0002 | | | | | | | | | |
| NLS-mEGFP | <0.0001 | 0.9999 | | | | | | | | |
| NLS-mNG | 0.0005 | 0.9989 | 0.9941 | | | | | | | |
| NLS-mKOkappa | <0.0001 | 0.4386 | 0.4153 | 0.0917 | | | | | | |
| NLS-stagRFP | <0.0001 | 0.0970 | 0.079 | 0.0123 | 0.9952 | | | | | |
| NLS-mScarlet | <0.0001 | 0.2822 | 0.2555 | 0.0500 | 0.9999 | 0.9998 | | | | |
| NLS-mCherry | <0.0001 | 0.9964 | 0.9979 | 0.7742 | 0.8465 | 0.3047 | 0.6707 | | | |
| NLS-emiRFP670 | 0.0513 | 0.4015 | 0.2636 | 0.7466 | 0.0039 | 0.0005 | 0.0019 | 0.0655 | | |
| NLS-miRFP670nano3 | 0.0002 | <0.0001 | <0.0001 | <0.0001 | <0.0001 | <0.0001 | <0.0001 | <0.0001 | <0.0001 | |
